# Haplotype-enhanced inference of somatic copy number profiles from single-cell transcriptomes

**DOI:** 10.1101/2022.02.07.479314

**Authors:** Teng Gao, Ruslan Soldatov, Hirak Sarkar, Adam Kurkiewicz, Evan Biederstedt, Po-Ru Loh, Peter Kharchenko

## Abstract

Genome instability and aberrant alterations of transcriptional programs both play important roles in cancer. However, their relationship and relative contribution to tumor evolution and therapy resistance are not well-understood. Single-cell RNA sequencing (scRNA-seq) has the potential to investigate both genetic and non-genetic sources of tumor heterogeneity in a single assay. Here we present a computational method, Numbat, that integrates haplotype information obtained from population-based phasing with allele and expression signals to enhance detection of CNVs from scRNA-seq data. To resolve tumor clonal architecture, Numbat exploits the evolutionary relationships between subclones to iteratively infer the single-cell copy number profiles and tumor clonal phylogeny. Analyzing 21 tumor samples composed of multiple myeloma, breast, and thyroid cancers, we show that Numbat can accurately reconstruct the tumor copy number profile and precisely identify malignant cells in the tumor microenvironment. We uncover additional subclonal complexity contributed by allele-specific alterations, and identify genetic subpopulations with transcriptional signatures relevant to tumor progression and therapy resistance. We hope that the increased power to characterize genomic aberrations and tumor subclonal phylogenies provided by Numbat will help delineate contributions of genetic and non-genetic mechanisms in cancer.

## Introduction

Copy number variations (CNVs) and loss of heterozygosity (LoH) events are major genome aberrations found in nearly all cancer cells. Characterization of CNVs in healthy and malignant tissues has informed the early detection, modes of progression, and resistance mechanisms of cancer^1–3^. However, the functional impacts of CNVs on the overall cellular activity, and how they drive malignant transformation remain largely unclear. In terms of clinical management, genome instability is also a key contributor to intratumoral heterogeneity. Therapy-resistant subclones frequently arising in the course of treatment pose a major challenge to cancer therapies. In addition to genetic heterogeneity, resistance may also stem from changes in the epigenetic or regulatory state, though the relative importance of different mechanisms has been difficult to establish^4^. All such changes, however, including genomic alterations, are likely reflected in the transcriptional state of the cell.

Single-cell RNA sequencing (scRNA-seq) methods provide an excellent opportunity to bridge genetic heterogeneity with the overall cellular state. It has been demonstrated that CNVs can be inferred from transcript abundance as well as allelic imbalance in heterozygous SNPs^5–8^. Reliable inference of copy number states, however, remains challenging using either approach due to sparse and noisy nature of single-cell measurements. Expression-based methods infer the presence of CNVs based on a general expectation that local amplifications or deletions will on average result in up- or down-regulation of genes within the affected region of the genome, respectively. Such approach can produce false-positive results due to local variations in expression unrelated to genomic copy numbers^9^. Allele-based approaches examine deviations of the relative allele frequency (“B-allele frequency” or BAF) caused by CNVs, and are less affected by sample or cell type variations^5,8^. They are hindered, however, by data sparsity and allele-specific transcriptional stochasticity in single cells^10^.

Existing approaches for CNV detection from scRNA-seq do not utilize the prior knowledge of haplotypes, or the individual-specific configuration of variant alleles on the two homologous chromosomes, which can enable more sensitive detection of allelic imbalance. Although current sequencing technologies are generally not haplotype-resolved, population-based phasing provides means to computationally infer the haplotype of an individual using patterns of linkage disequilibrium (LD) in the human population^11,12^. The estimated haplotypes are highly accurate within adjacent genomic regions, with a typical span of 50kb - 1Mb, but are subject to phase switch errors that accumulate with longer genomic distance. Nonetheless, population-based phasing has been successfully applied to characterize chromosomal aberrations in the context of germline polymorphisms as well as somatic evolution, mainly using DNA sequencing/array genotyping data^3,13–15^. The utility of phasing in detecting CNV signals from scRNA-based assays, however, has not been explored. We hypothesized that prior phasing information would be particularly impactful in the context of sparse coverage provided by scRNA-seq.

Finally, single-cell sequencing provides a unique opportunity to dissect genetically heterogeneous subpopulations, masked in traditional bulk measurements. Since scRNA-seq yields limited coverage per cell, methods that utilize allele information typically rely on aggregating information across cells (forming *in silico* pseudobulk profiles) to confidently define aberrations^5,8^. This approach, however, will only increase statistical power if the aberrations are shared between the cells included in the pseudobulk, and could lead to a dilution of signal with the inclusion of genetically distinct cells. Therefore, reliable identification of subclonal CNV events depends on the accurate inference of clonal cell populations at the same time.

We therefore developed a computational method, Numbat, which integrates expression, allele, and haplotype information derived from population-based phasing to comprehensively characterize the CNV landscape in single-cell transcriptomes. Numbat employs an iterative approach to jointly reconstruct the subclonal phylogeny and single-cell copy number profile of the tumor sample. Applying our method to 21 tumor samples (56905 single cells) representing a variety of cancer types and genomic complexity, we show that Numbat reconstructs high-fidelity copy number profiles from scRNA-data alone, and accurately distinguishes cancer cells from normal cells in the tumor microenvironment. Within heterogeneous tumors, Numbat readily identifies distinct subclonal lineages that harbor haplotype-specific alterations. Numbat does not require sample-matched DNA data or *a priori* genotyping, and is widely applicable to a wide range of experimental settings and cancer types.

## Results

### Enhanced detection of subclonal allelic imbalances using population-based haplotype phasing

Prior phasing information can effectively amplify weak allelic imbalance signals of individual SNPs induced by the CNV, by exposing joint behavior of entire haplotype sequences and thereby increasing the statistical power^3,15^. To examine the extent to which population-based haplotype phasing can be carried out from scRNA-seq data, we first analyzed a triple-negative breast cancer sample (TNBC4), containing wide-spread loss of heterozygosity. Using the reads from immune cells to select and genotype germline variants, we performed haplotype phasing using a reference-based phasing algorithm, Eagle2^11^. Based on the alleles that can be confidently phased using observed counts in the LoH regions (*P*<0.05, binomial test), we compared phasing performance with respect to two different population genome reference panels: TOPMed and 1000 genome (1000G)^16,17^.

We found that population-based phasing was effective at inferring the haplotype of long stretches of expressed SNPs (mean: 11.5 SNPs, IQR 2-16, TOPMed). SNPs within the same gene were phased with especially high accuracy (96.8%) as compared to co-expression based phasing (71.4%)^18^. Furthermore, population-based phasing was also able to infer the haplotype across multiple genes, producing perfectly phased blocks containing on average 3.5 genes (IQR 1-4) and achieving a between-gene phasing accuracy of 77.8%. In contrast, co-expression based phasing relies on haplotype-specific expression of alleles within the same gene and cannot phase across genes. The ability to infer phasing between genes is particularly useful for CNV inference, as it provides means to overcome stochastic allele-specific expression effects which give rise to bursts of gene-specific allelic imbalances in individual cells^5,19^. The differential phasing accuracy from within and between genes reflects the fact that the strength of genetic linkage decays with increasing distance (Supplementary Figure 1a).

Hidden Markov models have been effectively used to detect allelic imbalances from noisy signals^3,5,8,15,20,21^. The conventional allele-focused approach (haplotype-naïve HMM, such as that used by HoneyBADGER) infers the presence of events by the increased variance of allele frequencies in the affected regions (Figure 1a, first panel)^5,8,20,21^. On the other hand, a haplotype-aware HMM exploits signed deviations of phased haplotype frequencies to gain additional statistical power (Figure 1a, last panel)^3,15^. The aberrant genome state is represented by a pair of mirrored states with reciprocal transitions to account for phase switch errors in the population-derived haplotypes, which can shift between the more abundant haplotype (major haplotype) and the less abundant haplotype (minor haplotype, Supplementary Figure 1b). To reflect the decay in phasing strength over longer genetic distances, we introduced site-specific transition probabilities between haplotype states in the Numbat allele HMM (see Methods).

**Figure 1:**
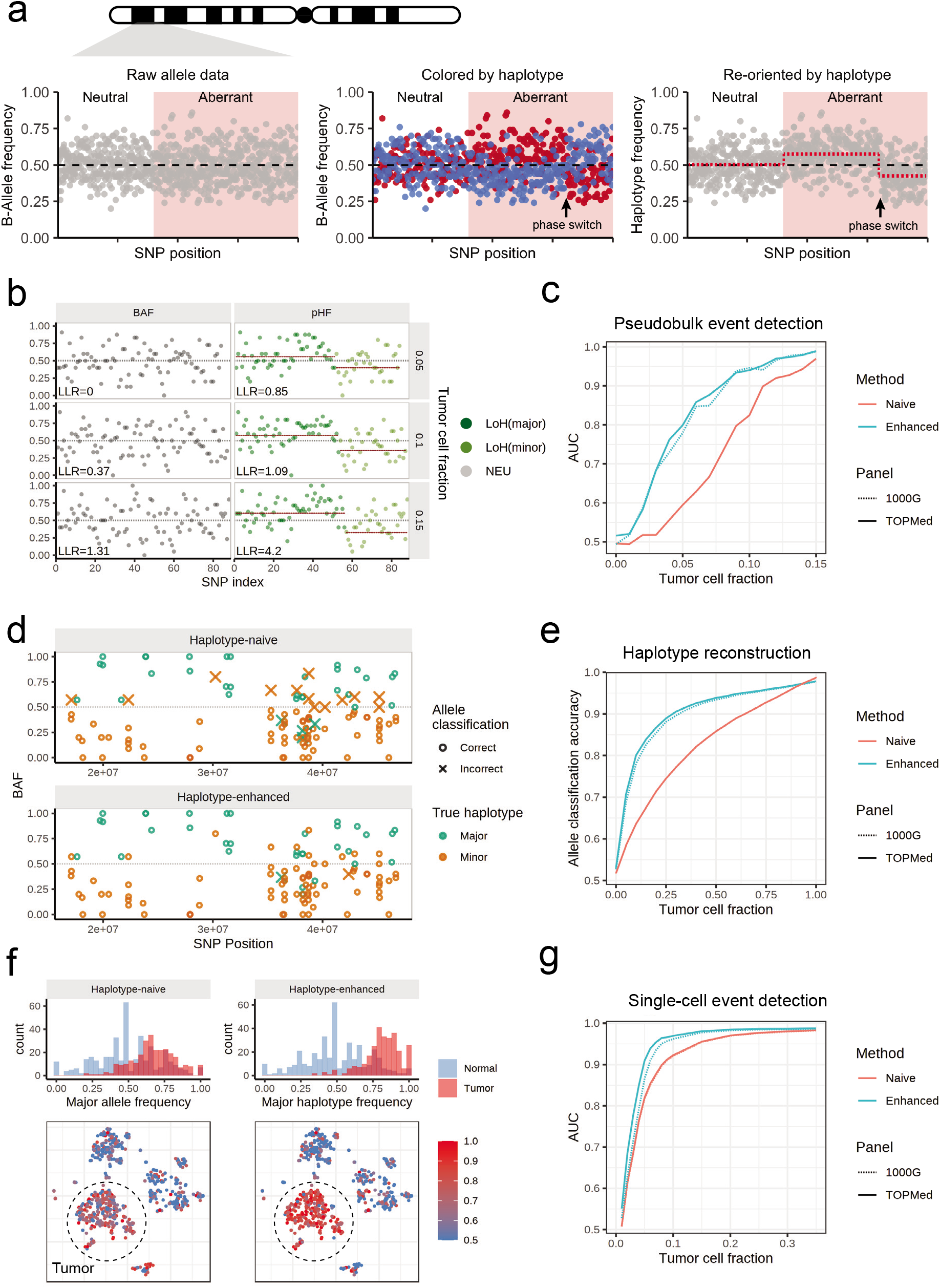
Population-based haplotype phasing enables sensitive detection of subclonal allelic imbalances in single-cell transcriptomes. **(a)** Schematic of using phasing signal to detect allelic imbalance. BAF, B-allele frequency. Simulated BAF signals are shown for a neutral and aberrant region harboring subclonal CNV. After BAF is transformed into haplotype frequency based on phase information, a Hidden Markov Model can be applied to detect aberrant regions and points of phase switches. **(b)** Example of statistical phasing signal uncovering subclonal allelic imbalances in TNBC4 *in silico* serial dilution experiment that are undetectable using BAF deviation. LLR, log-likelihood ratio. **(c)** Performance of allelic imbalance detection in simulated pseudobulks with (Enhanced) and without (Naive) haplotype phasing. AUC, area under the ROC curve. **(d)** Example of population phasing prior informing allele classification into major/minor haplotypes. **(e)** Performance of allele classification accuracy in simulated pseudobulks. **(f)** Example of population phasing prior improving detection of allelic imbalance in single cells. **(g)** Performance of allelic imbalance detection in single cells.

To benchmark the extent to which phasing helps with inferring CNVs and single-cell genotypes from scRNA-seq, we used the existing annotation of tumor and normal cells to create simulated pseudobulk profiles for a range of tumor cell fractions (clonality: 0-100%, Supplementary Figure 3). Compared to the naive model that only uses absolute deviations in allele frequency, the haplotype-enhanced allele HMM readily identified subtle allelic imbalances that would otherwise be invisible (Figure 1b) and achieved a higher AUC (by up to 21%) at low tumor fractions (1-15%) (Figure 1c). We then asked whether we can confidently test for the presence of individual events in single cells using the event characteristics obtained from the pseudobulk^5^. Accurately phased haplotype is crucial for identifying genotypes of individual cells, as it helps overcome the sparse SNP coverage by aggregating allele counts over affected regions^5^. In a naïve HMM, the assignment of alleles to either haplotype is solely based on the observed allele frequencies (an allele is classified as major if its BAF is higher than 0.5), whereas a haplotype-aware HMM combines evidence from prior phasing information and observed allele data to reconstruct haplotypes *a posteriori*. Using the BAF-based allele classification in the all-tumor pseudobulk as ground truth, we found that our haplotype-enhanced HMM achieved higher allele classification accuracy in aberrant regions especially at low tumor cell fractions (89% vs 75% accuracy at a 25% tumor cell fraction) (Figure 1d,e). As a result, allelic imbalances were more readily discernable in individual tumor cells using posterior haplotypes from the enhanced HMM (Figure 1f,g). Therefore, incorporating population phasing signal enables more sensitive characterization of allelic imbalances, and hence CNVs and LoH events, from scRNA-seq data.

### Accurate copy number inference from single-cell transcriptomes

Both the allelic imbalance, which reflects relative copy number of the two homologous chromosomes, as well as the changes in expression magnitudes, which reflect the total chromosomal dosage, provide signals for characterizing genome aberrations^5,8^. To integrate these two types of signals, we designed a joint HMM based on a generative statical framework (Methods, Supplementary Figure 2). We expanded the state space of the haplotype-aware allele HMM by combining the expected expression shifts and allelic ratio corresponding to each copy number configuration (Methods, Supplementary Figure 1c). To increase robustness, Numbat models gene expression as integer read counts using a discrete Poisson Lognormal mixture distribution, and accounts for excess variance in the allele frequency (e.g. due to allele-specific detection or transcriptional bursts) using a Beta-Binomial distribution. The Numbat HMM simultaneously calls significantly altered regions and determines their allele-specific copy number state and event boundaries, achieving joint copy number segmentation and inference (Figure 2a). The expression and allele signal in single cells can similarly be integrated via the generative model to produce probabilistic estimates of event presence in single cells (Figure 2b).

**Figure 2:**
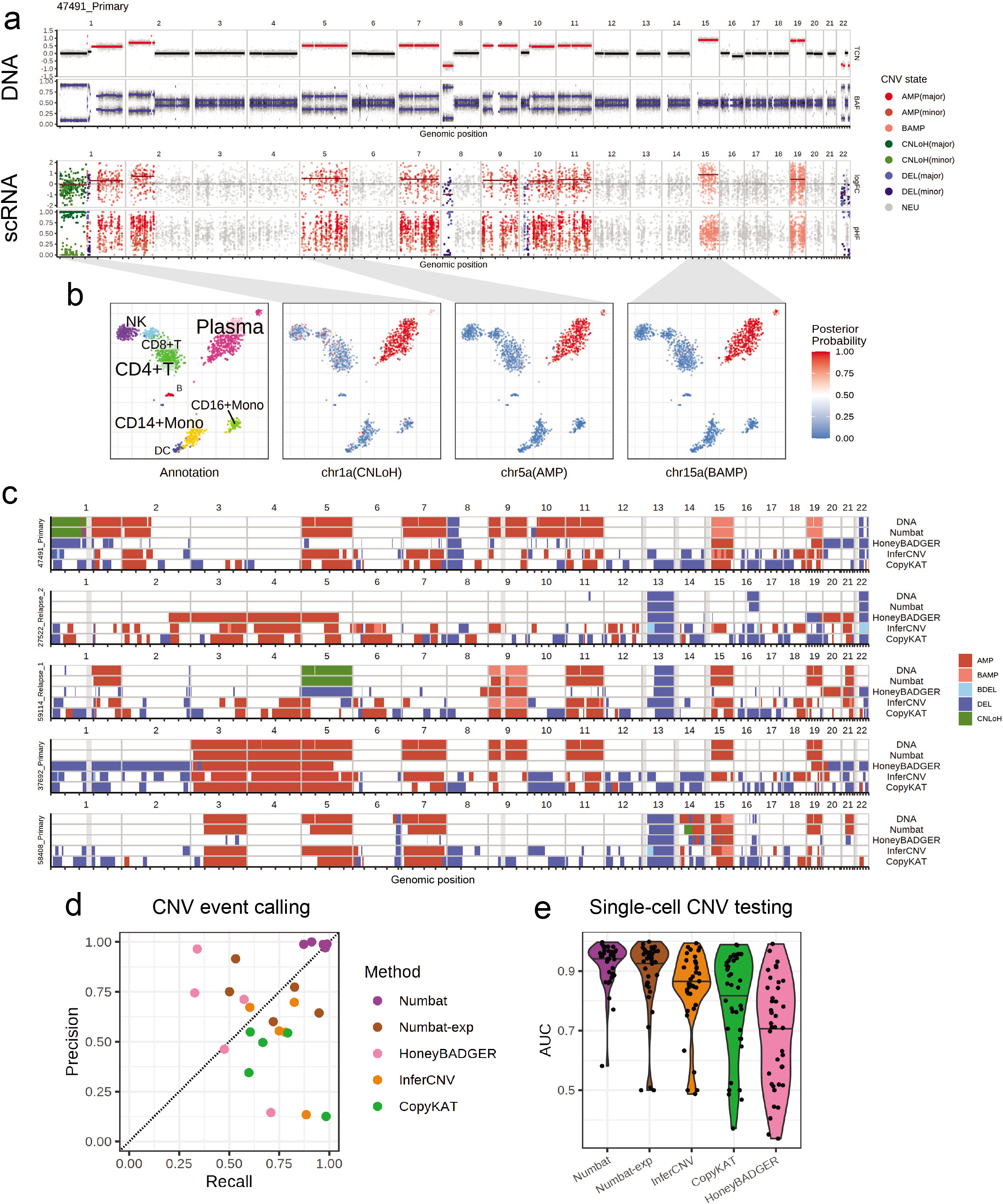
Numbat achieves accurate single-cell copy number inference via joint evaluation of gene expression, allele fraction, and prior haplotype phasing information. **(a)** Copy number profile of a primary multiple myeloma sample reconstructed by the joint HMM juxtaposed with the DNA profile. logFC, log fold-change. pHF, paternal haplotype frequency. TCN, total copy number. **(b)** Cell type annotation and posterior probability of CNV events in single cells visualized by a t-SNE embedding of gene expression profiles. **(c)** Copy number events detected by WGS, Numbat, and other methods. BAMP, balanced amplification. BDEL, balanced deletion. **(d)** Performance of CNV event detection by different methods. Each dot represents a distinct sample. **(e)** Performance of single-cell CNV testing by different methods. Each dot represents a distinct CNV event. Center line, median.

Existing methods infer copy number variations relative to the median ploidy, which can dilute signals of aberrant regions or mistake neutral regions for aberrant due to baseline shifts caused by hyperdiploidy or hypodiploidy^22^. To identify the diploid baseline, Numbat adopts a two-step approach: first, allelically balanced regions are identified through an allele-only HMM. The balanced regions are then clustered based on the expression shifts relative to the normal reference, and the cluster with the lowest average expression fold change is then designated as diploid regions (see Methods).

To validate the performance of copy number inference using Numbat, we first analyzed scRNA-seq data of 5 multiple myeloma (MM) samples with sample-matched, flow-sorted WGS (Supplementary Table 1). We detected CNV events from the malignant plasma cells using the Numbat joint HMM, Numbat expression-only HMM, and three other methods (HoneyBADGER, InferCNV and CopyKat). We found that the copy number events identified by Numbat are highly concordant with the corresponding DNA profiles (Figure 2c, Supplementary Figure 4), achieving higher overall accuracy (precision: 98.7%, recall: 94.5%) than other methods (Figure 2d). Compared to the expression-only HMM, incorporating allele information significantly improved the event calling performance. In addition, Numbat correctly identified copy-neutral loss of heterozygosity (CNLoH) events in two samples (chr1p of 47491_Primary, chr5 of 59114_Relapse_1), which are invisible to approaches that consider only expression magnitude, including InferCNV and CopyKAT (Figure 2c). Numbat also out-performed other methods on CNV testing (given known event boundaries) on a single-cell level (Figure 2e).

Importantly, Numbat correctly identified the diploid baseline in all 5 cases, whereas the copy number estimates produced by the other three methods are often confounded by baseline shifts caused by hyperdiploidy (e.g. 37692_Primary and 47491_Primary, Figure 2c). This issue is particularly pronounced in a pre-malignant breast cancer sample (DSCI1), where CopyKAT denoted the majority of chromosomes 3,9,10,15 as deleted, and chromosomes 1,7,14 as copy-neutral (Supplementary Figure 5a). In contrast, Numbat analysis using both allele and expression data revealed that chromosomes 3,9,10,15 are allelically balanced and therefore likely remain in diploid state, whereas chromosomes 1,7,14 carry wide-spread allelic imbalance around ⅔ fraction and are likely in triploid state (Supplementary Figure 5b,c).

### Iterative strategy to decompose tumor clonal architecture

scRNA-seq is commonly used to examine a full spectrum of cell states within the tumor microenvironment, including different malignant, immune and stromal subpopulations, whose classification is often unknown in advance. Therefore, reconstructing the single-cell copy number aberrations in heterogenous cell populations requires the inference of clonal populations and genomic aberrations at the same time. In heterogenous tumors, cells with distinct genotypes can generally be assumed to have originated from a common cell of origin, and are thus related to each other via a phylogeny. Their evolutionary relationships, if known, can be exploited to improve CNV detection by sharing information across cells in the same lineage^23^. On the other hand, given an estimated single-cell copy number profile, a CNV-based tumor phylogeny can be inferred^24,25^. To perform joint inference of single-cell CNV profiles and the associated subclonal phylogeny, we have adopted an alternating optimization procedure. In each iteration, Numbat first identifies CNVs in each branch of clonal phylogeny using the joint HMM on pseudobulk expression and allele profiles (Figure 3a). It then evaluates the evidence for each unique event in individual cells using a Bayesian hierarchical model, producing a matrix of posterior probabilities of CNVs by cell (Figure 3b). Next, to recover the tumor clonal architecture, Numbat employs a maximum-likelihood perfect phylogeny approach to fully propagate the uncertainty in single-cell CNV calls^26^. Briefly, the genotype probabilities are used to search for an optimal tree topology using nearest neighbor interchange (NNI), and mutations are placed on the tree based on maximum likelihood (Figure 3c). Numbat then uses the refined single-cell phylogeny to form more precise lineage-specific pseudobulks, iteratively optimizing single-cell copy number profiles and tumor phylogeny. The iterative approach can be initialized in different ways. By default, Numbat approximates an initial phylogeny by hierarchical clustering of window-smoothed expression signals. Alternative initialization strategies can include, for example, a hierarchy of expression-based cell subpopulations.

**Figure 3:**
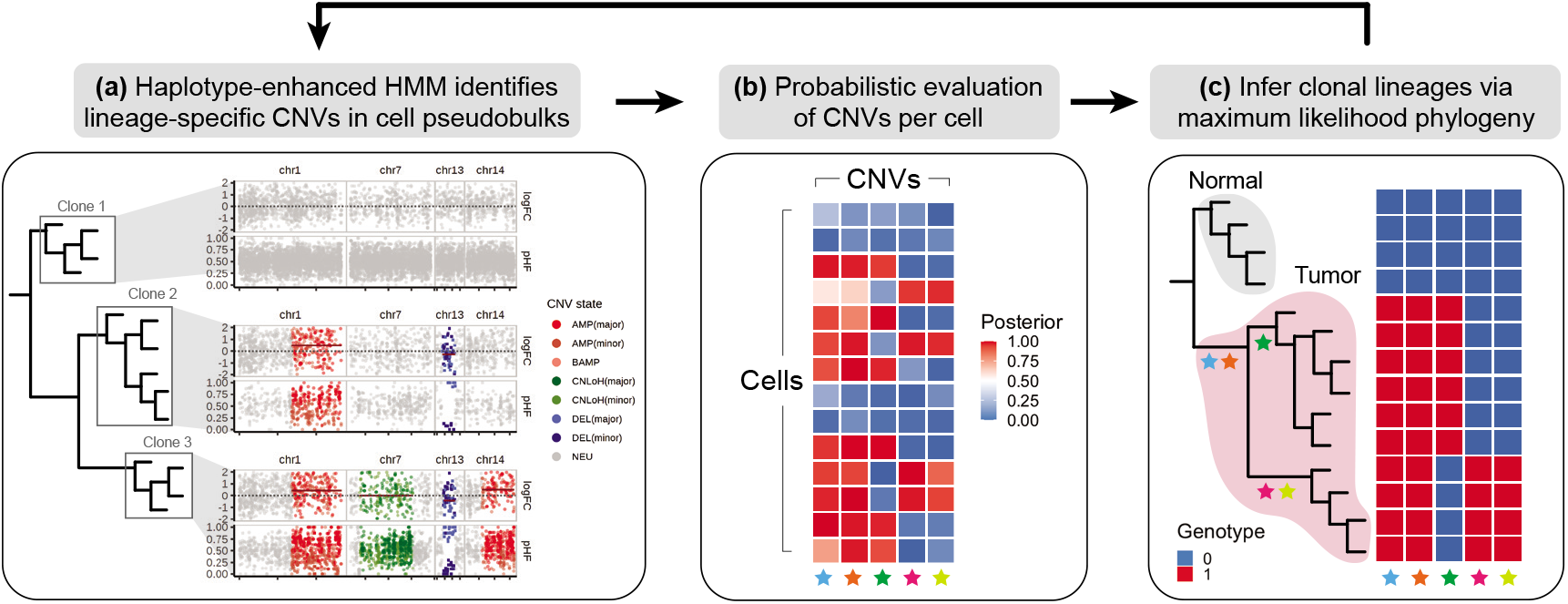
Iterative strategy to identify tumor subclones. **(a)** Numbat groups cells into pseudobulk profiles by major clades in the single-cell phylogeny, and runs a haplotype-enhanced HMM on each pseudobulk to identify lineage-specific CNVs. **(b)** Numbat evaluates the presence/absence of each CNV in each cell probabilistically using a Bayesian hierarchical model. **(c)** Numbat then infers a maximum-likelihood phylogeny that captures the evolutionary relationships between single cells.

### Reliable identification of cancer cells in the tumor microenvironment

Precisely distinguishing the malignant cells within heterogeneous cell mixtures is a well-established problem^6,9^. Since the non-malignant cells do not share aberrations with the tumor, the tumor population should be isolated as a distinct clade in the reconstructed clonal phylogeny (Figure 3c). To systematically benchmark Numbat’s ability to distinguish tumor cells from non-malignant cells in the tumor microenvironment, we analyzed 5 triple-negative breast cancer (TNBC) samples and 5 anaplastic thyroid cancer (ATC) samples in addition to 8 MM samples (Supplementary Table 1). We defined true tumor cell clusters based on the expression of well-established cell-type or tumor-specific markers (*EPCAM* for TNBC^27^, *KRT8* for ATC^28^, *MZB1* for MM) as well as aneuploidy status (Methods). The overall tumor versus normal cell classification accuracy of Numbat was consistently higher than that of CopyKAT in the three tumor series (Supplementary Figure 6). The average classification accuracy for Numbat was 98.5% on TNBC and 98.7% on ATC series, whereas CopyKAT achieved an average accuracy of 98.1% on TNBC and 98.5% on ATC series (Supplementary Figure 6–8; ATC5 sample was excluded from benchmark due to the lack of clear expression of tumor marker *KRT8*). On the 8 MM samples, we found that Numbat maintained a stable performance (98.8%) whereas CopyKAT misclassified clusters of cells in five out of eight samples (Supplementary Figure 9), resulting in lower performance (74.8%, Supplementary Figure 6). The reduced accuracy of CopyKAT in the MM series is likely due to the lower sequencing coverage per cell and the less pronounced chromosomal aberrations in those samples. Numbat integrates two orthogonal lines of evidence (expression and allele) for aneuploidy status in each cell, enhancing signal and reducing the possibility of deriving erroneous conclusions from either source of information alone (Supplementary Figure 6–9).

### Allele-specific CNV analysis reveals additional subclonal complexity

Application of the Numbat iterative strategy to TNBC and ATC datasets also identified subclonal tumor populations in multiple samples (Figure 4, Supplementary Figure 10). In particular, we found that allelic imbalances frequently contributed to the clonal complexity of tumors. For example, in TNBC1, Numbat inferred a branching phylogeny composed of two major subclonal lineages undergoing concurrent evolution (Figure 4a). The two lineages share CNLoH events on multiple chromosomes (i.e. chr1p, chr13, chr14, chr17, chr19, Figure 4a). Numbat also identified subclonal CNLoH events on chr3p and chr22q that are exclusive to the minor lineage (Figure 4b,c). Such copy-neutral events do not exhibit relative deviations in expression magnitude and can only be identified through allele analysis (Supplementary Figure 11). In addition, Numbat revealed that the major lineage carries an imbalanced amplification on chr16 whereas the minor lineage carries an allelically-balanced amplification on the same chromosome. Although both lineages carry an amplification on chr15 with similar increase in expression magnitudes (Figure 4b), their haplotype frequencies appear to be mirrored (Figure 4d), indicating that different homologous copies of the chromosome were duplicated independently in the evolutionary history of the two clones (Figure 4e). Another example of an unusual clonal divergence pattern can be seen in ATC1. While the overall expression profile suggested that ATC1 harbors a relatively simple genome (Supplementary Figure 11), Numbat’s analysis revealed two diverging tumor lineages with reciprocal aberrations. While one subclone harbors an amplification on chr7 and a CNLoH on chr17, the other harbors a CNLoH on chr7 and an amplification on chr17 (Figure 4f-i). Recent studies using scDNA-seq data revealed that such multi-allelic and mirrored CNVs are prevalent sources of tumor heterogeneity^29,30^. These events, however, have not been previously inferred from scRNA-seq due to limited resolution in allele analysis and the lack of signal in the overall expression profile. In summary, these examples illustrate that the integration of phased allele data with expression signals can aid in the detection of subclonal events and phylogenies reflecting dynamic clonal complexity of evolving tumors.

**Figure 4:**
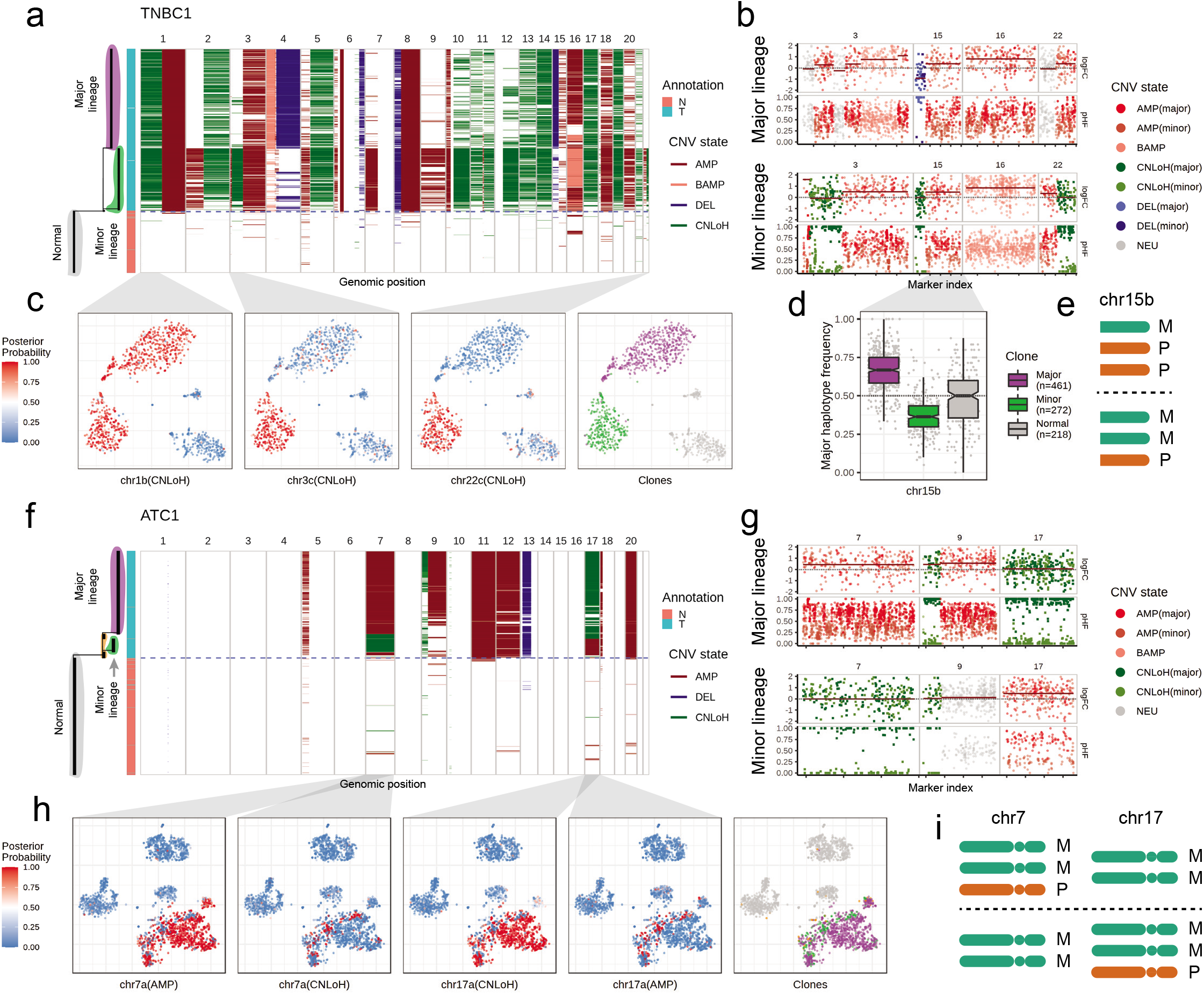
Numbat reveals additional complexity in tumor subclones through allele-specific copy number analysis. **(a)** Single-cell CNV landscape and reconstructed phylogeny of TNBC1. Branch lengths correspond to the number of mutations. Blue dashed line separates predicated tumor and normal cells. The annotation corresponds to cell type ground truth (N, normal. T, tumor). **(b)** Bulk CNV profile of the major and minor lineage. pHF, paternal haplotype frequency. **(c)** Posterior CNV probability of shared and lineage-sepcific CNVs in a t-SNE embedding of gene expression profiles. **(d)** Major haplotype frequency in single cells. Only cells with at least 5 total allele counts in the region are shown. Center line, median; box limits, upper and lower quartiles; whiskers, 1.5x interquartile range. **(e)** Schematic of copy number state of chr15b in the major and minor lineage. M, maternal. P, paternal. The designation of maternal and paternal chromosomes is arbitrary. **(f)** Single-cell CNV landscape and reconstructed phylogeny of ATC1. **(g)** Bulk CNV profile of the major and minor lineage. **(h)** Posterior CNV probability of subclonal multi-allelic CNVs in a t-SNE embedding of gene expression profiles. **(i)** Schematic of copy number state of chr7 and chr17 in the major (top) and minor (bottom) lineages.

### Unraveling the interplay between genetic and transcriptional heterogeneity in tumor evolution

The decomposition of genetic subclones from single-cell transcriptomes provides an opportunity to jointly characterize genetic and transcriptional heterogeneity during the course of tumor evolution. In particular, acquired copy number alterations can be used as natural genetic barcodes in conjunction with characteristic expression signatures to track the behaviors of clonal populations across time. We therefore applied Numbat to investigate the clonal evolutionary history of a therapy-resistant multiple myeloma (Patient 27522) with four sequential samples (primary, remission, first relapse, second relapse). Numbat identified three malignant subclones (g1-g3): one that harbors only ancestral deletions on chromosomes 13 and 22 (g1), one that harbors an additional chr1p deletion (g2), and one that has acquired a chr16q deletion (g3) (Figure 5a-c). At primary diagnosis, the tumor was only composed of clones g1 and g2, both of which appeared to be undetectable at the time of remission. However, clone g1 survived the therapy and reappeared at the first relapse. Furthermore, clone g1 also gave rise to clone g3, which continued to expand during subsequent therapy, and became the dominant tumor subclone at the second relapse (Figure 5c).

**Figure 5:**
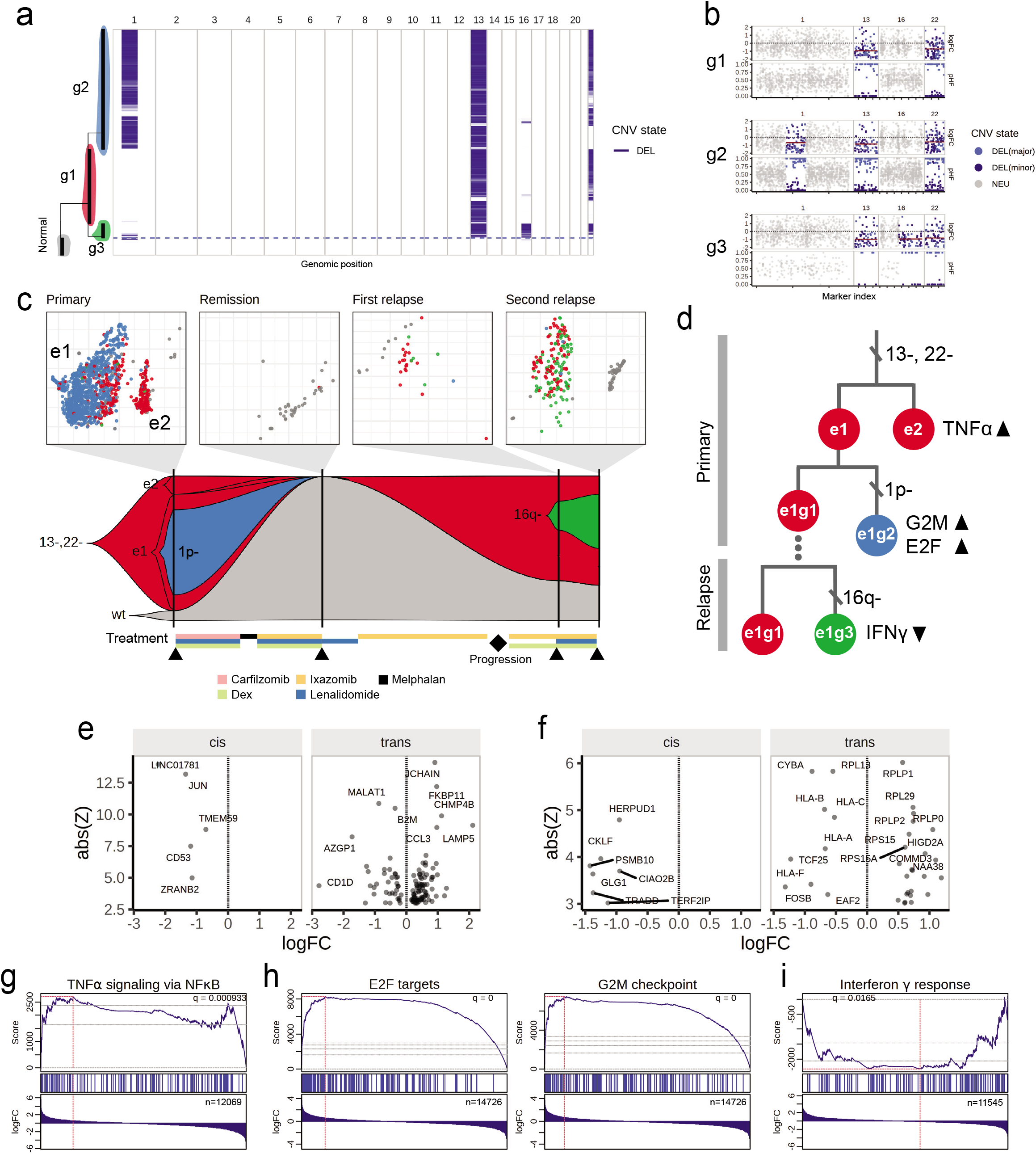
Tracking clonal evolution history of a therapy-resistant multiple myeloma using Numbat. **(a)** Integrated single-cell CNV landscape and phylogeny of plasma cells from all four samples. Only plasma cells are included. **(b)** Bulk CNV profile of three main tumor clones. **(c)** Clonal evolutionary history integrating genetic and transcriptional heterogeneity. Top, t-SNE embedding of gene expression profiles colored by genetic clones. The embeddings are created separately for each sample. Only cells with >90% posterior classification confidence are shown. Bottom, change in tumor clonal composition over time. At each time point, only clones with more than 5% cellular fraction are shown. **(d)** Genetic and transcriptional changes in the proposed evolutionary history. **(e)** Differentially expressed genes between e1g1 and e1g2 cells. **(f)** Differentially expressed genes between e1g1 and e1g3 cells. **(g)** Enriched pathway in e2g1 cells relative to e1g1 cells. **(h)** Enriched pathways in e1g2 cells relative to e1g1 cells. **(i)** Enriched pathway in e1g3 cells relative to e1g1 cells.

The tumor cells in the primary sample separate into two distinct transcriptional expression clusters (e1 and e2). While the ancestral clone g1 is found in both e1 and e2 subpopulations, the derived clone g2 appears to be restricted to cluster e1 (Figure 5c). This suggests that a large-scale shift in the transcriptional landscape gave rise to the two distinct tumor subpopulations (e1 and e2), which predated the chr1p deletion event within e1, or that with the acquisition of chr1p deletion, g2 tumor cells lost the ability to enter state e1 (Figure 5d). Integrating both aspects of heterogeneity, we resolved three main subpopulations in the primary sample: cells in expression cluster 1 with wildtype chr1 (e1g1), cells in expression cluster 1 with chr1p deletion (e1g2), and cells in expression cluster 2 (e2g1). Since clone g1 was the major cell population that re-emerged after remission, we asked whether it was derived from e1g1 or e2g1 cells. The g1 cells in the relapse sample carried the expression signatures of e1 cells from the primary sample, as evidenced by the shared differentially expressed genes (Supplementary Figure 12), indicating that the relapsed tumor likely originated from e1g1 cells in the primary sample (Figure 5d).

We next investigated the interplay between genetic and transcriptional subpopulations using differential expression and pathway enrichment analysis, separating likely *cis* and *trans* effects. Comparing e1 and e2 cells with the same copy number background (e2g1 vs e1g1) in the primary tumor, we found that e2 cells have higher activation of the tumor necrosis factor α (TNFα) signaling pathway (Figure 5g, Supplementary Table 2). It has been shown that TNFα triggers the release of IL-6, a myeloma growth factor, by activating nuclear factor kappa B (NF_K_B)^31^. Comparing e1 cells with and without the chr1p deletion (e1g1 vs e1g2), we found that cells with chr1p deletion have higher activation of pathways associated with cell cycle (G2M checkpoint and E2F targets), indicating a hyper-proliferative phenotype (Figure 5h, Supplementary Table 2). Differential gene expression analysis between e1g1 and e1g2 cells revealed 5 significantly differentially expressed genes in *cis* of the chr1p deletion event (i.e. in the affected genomic region) and 117 genes in *trans* (i.e. outside of the affected genomic region, Figure 5e). All 5 DE genes in *cis* of the deletion are significantly down-regulated, as expected. The genes involved in the enriched pathways do not overlap significantly with the deleted region (*P*=1, E2F targets; *P*=1, G2M checkpoint; two-sided binomial test), indicating that those transcriptional changes may be driven by processes other than the CNVs we have detected. The two genetic subclones in the second relapse sample (g1 and g3) do not separate into distinct expression clusters (Figure 5c). Direct comparison of their expression patterns, however, revealed 7 significantly differentially expressed genes in *cis* and 33 in *trans* of the deletion (Figure 5f), and showed that the cells carrying chr16q deletion have significantly downregulated interferon gamma (IFNγ) response pathway (Figure 5i, Supplementary Table 2). Similar to the previous case, the genes involved in the enriched pathways do not overlap significantly with the deleted region (*P*=0.31, two-sided binomial test). IFNγ signaling plays an important role in tumor cell clearance by immune surveillance, and its dysregulation is associated with immune evasion and poor response to immunotherapy^32^. This is consistent with the more aggressive phenotype of clone g3, which achieved clonal dominance after several rounds of therapy (Figure 5c).

## Discussion

Tumor plasticity and the resulting therapy resistance can be driven by both genetic and non-genetic mechanisms, such as large-scale chromatin remodeling or aberrant activation of transcriptional programs^4,33^. The interplay between the genetic and non-genetic mechanisms and their relative importance remain poorly understood. Methods that can reliably infer genetic alterations from a cell’s transcriptome have the potential to illuminate these effects by characterizing both aspects of intratumoral heterogeneity at single-cell resolution.

Compared to DNA-based approaches, scRNA-seq provides limited coverage of alleles and additionally suffers from transcriptional noise. Numbat attempts to address those challenges by incorporating prior haplotype information obtained from population-based phasing. We show that prior phasing signals can be integrated with allele and expression information in a Hidden Markov Model to enhance detection of subclonal copy number alterations from scRNA-seq data. The power of the haplotype-enhanced HMM included in Numbat can be further improved by more accurate haplotype information obtained from other techniques such as long-range haplotype phasing that takes advantage of individual relatedness^34^ or experimental approaches that resolve haplotypes^35^.

Reconstructing the single-cell copy number profile from heterogenous cell populations requires the inference of clonal populations and genomic aberrations at the same time. Aberrations are more evident when aggregating information across cells, yet not all of the cells may share any given event, setting up a conundrum. Numbat solves this problem by iteratively inferring the tumor phylogeny using detected aberrations and refining single-cell copy number estimates by exploiting the structure of the tumor phylogeny. Application to three tumor series (ATC, TNBC, MM) showed that Numbat precisely differentiated normal and malignant cells (marked by aneuploidy) in the tumor microevironment and revealed additional subclonality within the tumor population. However, Numbat shares the same limitation with existing methods in that determining the number of confident subclones still relies on manual inspection of the tumor phylogeny and copy number profile^5–8^. In addition, since Numbat detects events from cell pseudobulks, it is possible that CNVs carried by rare cell populations are missed. These rare subclones may carry sufficient evidence to establish that some additional aberrations are present, but not to define the aberrations themselves (e.g. boundaries, haplotype configuration) with sufficient confidence.

Uncertainties in CNV inference from scRNA-seq can stem from a variety of factors, including local variations in cell-type specific expression, transcriptional stochasticity, and coverage variation in individual cells. Therefore, it is crucial to quantify the uncertainty in the copy number estimates for downstream analysis. By adopting a Bayesian statistical framework and performing cell-specific noise profiling, Numbat provides a natural way to integrate evidence from different sources of signals as well as to derive confidence estimates for CNV events and clonal assignments.

Tumor baseline ploidy estimation is a challenging problem in copy number analysis^22,36^. Current methods, including those that integrate allelic information, infer copy number variations relative to the median ploidy, which can be confounded by hyperdiploidy or hypodiploidy^22^. Numbat attempts to address this problem by adopting a strategy previously developed in DNA analysis^22,37^. This approach was effective and correctly identified diploid regions in 5 tumor samples with WGS validation. However, challenges remain in cases when shared neutral regions are scarce (e.g. TNBC1) or when allele coverage is poor, in which case manual curation is still needed. Further improvements will be needed to robustly determine copy number baseline in tumors with complex copy number profiles.

Allele-specific CNV analysis has shown major advantages over total copy number analysis in studies of cancer genomes^29,30,38^. Although variations in chromosomal dosage are often discernable from large-scale gene expression changes, CNLoH events and haplotype-specific alterations can only be detected using allele information. Numbat analysis of previously published tumor samples revealed additional subclonal complexity resulting from haplotype-specific alterations, highlighting the importance of allele-specific copy number analysis. Finally, to demonstrate the type of integrative analysis enabled by Numbat, we used it to characterize the genetic and transcriptional subpopulations in a serial multiple myeloma sample. Comparing the gene expression patterns of tumor subclones revealed that many of the transcriptional changes relevant to cancer progression and therapy resistance are likely independent of the copy number alterations that we detect, possibly due to epigenetic alterations or other genetic events (such as SVs, SNVs and Indels). Additional methods integrating genetic and epigenetic information will be needed to fully resolve the impact of genome instability on tumor cell states^29^. Meanwhile, we hope Numbat will serve as a valuable tool for the analysis of genetic and functional states in cancers, and facilitate development of improved computational approaches in this area.

## Methods

### Datasets

We analyzed 3 tumor series from two published studies (WASHU and MDA, Supplementary Table 1)^6,39^. The MM series from the WASHU dataset contains 10 samples of 5 multiple myeloma patients. We analyzed a subset of samples from the WASHU study because of the availability of high-quality sample-matched flow-sorted WGS.

### Pre-processing of scRNA-seq data

We used the Cell Ranger (version 6.0.2, 10x Genomics) software suite to process the raw FASTQ or BAM files obtained from the previously published studies. We used conos^40^ to perform multi-sample integration, clustering, and generation of graph embeddings.

### Genotyping and phasing from scRNA-seq data

To identify heterozygous and homozygous germline SNPs, we use cellsnp-lite^41^ to generate allele counts for a panel of known common SNPs (population allele frequency > 5%). SNPs with variant allele frequency (aggregating all cells) between 0.1 and 0.9 are identified as heterozygous. SNPs with ≥ 10 reads covering the alternate allele with VAF = 1 are identified as homozygous. We then use Eagle2^11^ to phase the identified heterozygous SNPs using established population reference panels (e.g. 1000 Genome and TOPMed).

### Statistical modeling of expression magnitude and allele frequency

We formulate a generative model for the observed read counts per transcript and the observed allele counts per SNP (Supplementary Figure 2). This model generalizes to both pseudobulk and single-cell setting. We aim to infer the DNA state for each marker, denoted as *g* = (*c*_*m*_, *c*_*p*_) where *c*_*m*_ is the number of maternal copies and *c*_*p*_ is the number of paternal copies. Note that in single cells, *c*_*m*_ and *c*_*p*_ can take any non-negative integer value. For example, in diploid regions, g = (1,1) whereas in a heterozygous loss of the paternal chromosome, g = (1,0). Since a pseudobulk can contain a mixture of cells in diploid state and cells in altered state, c_m_ and *c*_*p*_ can take any continuous value in the positive domain. For convenience, we reparameterize (c_m_, c_p_) as the change in total chromosome dosage relative to the diploid state (*ϕ*) and haplotype fraction (θ) as follows:

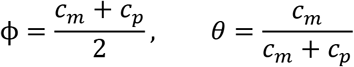

which are the targets of inference. Note that in single cells, ϕ and θ take on discrete values. In pseudobulks, ϕ ∈ [0, ∞) and θ ∈ [0,1], which depend both on the mixture proportion and the underlying genotype.

We observe two types of markers: expression counts per gene and allele counts per SNP. Gene expression counts are only emitted once per gene whereas allele counts are emitted at each SNP. For gene *i*, we denote the gene expression count as *X*_*i*_ . Note that both *X*_*i*_ and library size *l* are observed (obtained from upstream transcriptome quantification). For allele data, we use *Y*_*j*_ to denote the observed variant allele count of the *j*th SNP, and d_j_ to denote the total allele count (sum of reference allele count and variant allele count). Once the variant alleles are phased, *Y*_*j*_ is the maternal allele count. Given the framework, we model the gene expression count as a Poisson distribution where the rate parameter follows a Lognormal distribution:

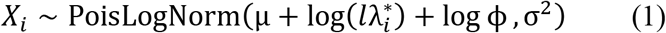

Here, 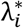 is the empirically estimated mean expression for gene *i* in the reference profile. Shared between all genes, μ and σ^2^ are hyperparameters representing the bias and variance in the log expression fold change between the reference and observation. We model maternal allele count for SNP *j* using a Beta-Binomial distribution:

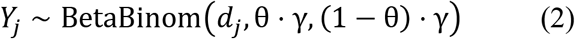

Where γ is a hyperparameter that accounts for the excess variance in allele counts caused by haplotype-specific expression.

### Phase switch probabilities

We modeled the occurrence of phase switch errors from population-based haplotype phasing along the genome using a Poisson process with a uniform rate λ. Between two adjacent SNPs with genetic distance (in centimorgan) *d*, the number of phase switches *W* can be modeled by a Poisson distribution:

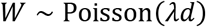

The probability of two SNPs being discordant in phase is therefore

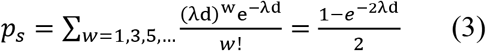

### Hidden Markov model to detect allele-specific copy number alterations

We designed an HMM that efficiently integrates expression deviation and haplotype imbalance signals to detect CNVs in cell population pseudobulks. The hidden states correspond to 15 possible copy number states depending on the variation in total copy number, extent of allelic imbalance, and haplotype state (major or minor, accounting for random phase switch errors in the population-derived haplotype; subclonal and clonal, accounting for the extent of imbalance).

| State ID | Copy number state | Clonal fraction | Haplotype state | $\phi$ | $\theta$ |
| --- | --- | --- | --- | --- | --- |
| 1 | Neutral | - | - | 1 | 0.5 |
| 2 | CNLoH | Subclonal | Major | 1 | $0.5 + \theta_{min}$ |
| 3 | CNLoH | Subclonal | Minor | 1 | $0.5 - \theta_{min}$ |
| 4 | CNLoH | Clonal | Major | 1 | 0.9 |
| 5 | CNLoH | Clonal | Minor | 1 | 0.1 |
| 6 | Amplification | Subclonal | Major | $1 + \phi_{min}$ | $0.5 + \theta_{min}$ |
| 7 | Amplification | Subclonal | Minor | $1 + \phi_{min}$ | $0.5 - \theta_{min}$ |
| 8 | Amplification | Clonal | Major | 2.5 | 0.9 |
| 9 | Amplification | Clonal | Minor | 2.5 | 0.1 |
| 10 | Deletion | Subclonal | Major | $1 - \phi_{min}$ | $0.5 + \theta_{min}$ |
| 11 | Deletion | Subclonal | Minor | $1 - \phi_{min}$ | $0.5 - \theta_{min}$ |
| 12 | Deletion | Clonal | Major | 0.5 | 0.9 |
| 13 | Deletion | Clonal | Minor | 0.5 | 0.1 |
| 14 | Balanced amplification | - | - | $1 + \phi_{min}$ | 0.5 |
| 15 | Homozygous deletion | - | - | $1 - \phi_{min}$ | 0.5 |

Each of the 15 hidden states described above emits a gene read count *X*_*i*_ and a maternal allele count *Y*_*j*_ according to the distributions specified by the equations (1) and (2), respectively. The transition probabilities are specified by *t* and *p*_*s*_, where *t* is the transition probability between copy number states, and *p*_*s*_ is the transition probability between haplotype states (i.e. phase switch probability between major and minor haplotypes). *t* is homogeneous in the Markov chain whereas *p*_*s*_ is site-specific. To reflect LD decay over genetic distance, we model *p*_*s*_ as a monotonically increasing function of genetic distance from the previous SNP according to equation (3).

To infer the hidden copy number states, we use the Viterbi algorithm to identify the most probable copy number states for each marker position. Although contiguous genomic segments can occupy distinct copy number states, which cannot be captured by any single set of *ϕ* and *θ*, we use one set of minimum-threshold parameters (*ϕ*_*min*_ and *θ*_*min*_) to initially identify all detectable CNVs with various deviation magnitudes. By default, we set *ϕ*_*min*_ = 1.2 and *θ*_*min*_ = 0.08. The true underlying dosage ratio and haplotype frequency are event-specific and are estimated separately by maximizing the total model likelihood. Finally, we obtain the haplotype classification of major/minor alleles based on the posterior marginal of each SNP, computed from the forward-backward algorithm using the maximum likelihood estimates of (ϕ, θ).

The allele-only HMM and the expression-only HMM are special cases of the joint HMM defined above. The allele-only HMM includes a subset of the states (1-5) and only allele counts are observed, whereas the expression-only HMM only uses the gene expression counts and does not allow transition between haplotype states.

### Identifying the diploid state

We use a graph-based clustering approach to identify genomic regions in the diploid (neutral) state from a given pseudobulk profile. First, regions of allelic imbalance are identified by the allele-only HMM and excluded. The remaining allelically balanced segments are assumed to be in even-valued copy number states^22^. We then perform a pairwise comparison of the log expression fold-change (logFC) of the balanced segments using Student’s t-test. We construct a graph where the nodes represent the balanced segments and the edges are determined by the obtained adjusted significance scores from the previous step. An edge connecting two segments means that their expression magnitudes are not significantly different, and that they likely occupy the same total copy number state. We note that a clique in such a graph would be a collection of segments that are most coherent in expression magnitude. Consequently, the diploid segments can be inferred as the maximum clique with the lowest average logFC^22^. An extension of this procedure can be used to identify shared diploid states in multiple pseudobulk profiles containing different cells in the tumor, where segments are compared in each pseudobulk and a Simes’ test is used to determine edges.

### Obtaining consensus CNV events from multiple pseudobulks

To resolve CNV calls from HMM on multiple pseudobulks into unique events, we apply the following heuristics. First, each called CNV is represented as a node in a graph, and an edge is added between pairs of nodes if the two CNV segments significantly overlap (length of the overlap is more than 45% of both segments). The nodes are then clustered by determining connected components of the resulting graph. Within each component, the CNVs are ranked by likelihood evidence (combined LLR of expression and allele deviation). All CNVs from the pseudobulk profile that harbored the top event is kept. For instance, let A and B be events called in pseudobulk 1 and C be an event called in pseudobulk 2, and C overlaps both A and B. If event B is the top event with the highest LLR, we include both A and B in the consensus profile.

### Single-cell CNV evaluation

We make inferences on the underlying genotype of individual cells jointly using the observed expression and allele counts. First, using the diploid regions identified in pseudobulk HMM, we estimate the cell-specific expression fold-change bias and variance by maximum likelihood:

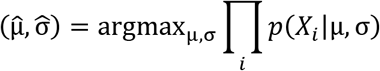

In a given genomic region of a given cell, the posterior probability of each genotype is obtained by

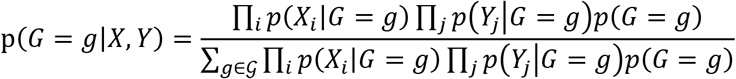

where the likelihood functions are defined according to the generative model described above. Although the maternal and paternal copy number can take any non-negative integer value, in practice we only consider seven possible genotypes: 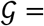 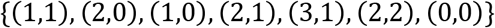.

### Maximum-likelihood phylogeny inference using uncertain genotypes

We implement a modified version of a recently described approach to infer a maximum-likelihood perfect phylogeny based on uncertain genotypes^26^. Using the CNV by cell genotype probabilities obtained in the previous step, we first construct an initial tree using the neighbour joining algorithm, where the distance between two cells is defined as the Euclidean distance of CNV posteriors:

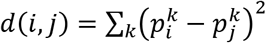

In large datasets, the initial tree can be constructed by the UPGMA algorithm instead to save computational time. We then search for an optimal tree topology that maximizes the genotype likelihood (as defined in ^26^) using the nearest neighbor interchange (NNI) algorithm. After identifying the optimal mutation placement on the tree by maximum likelihood, we simplify the mutational history (i.e. reduce the number of evolutionary steps) by re-assigning a mutation to the same branch as another mutation that happened directly up- or downstream. The cost for such re-assignment is defined by the corresponding decrease in the genotype likelihood. We iteratively perform the least costly reassignment until a specified maximum threshold is reached. The final phylogeny specifies not only the lineage relationship between cells but also the genotypes of each subclone (defined as a paraphyletic monophyletic group in the phylogeny sharing the same mutation profile). The tumor lineage can be identified as the clade in the phylogeny with the highest mutation burden (total number of mutations across all cells in the clade).

### Posterior assignment of cells to copy number profiles and clades

Given *K* genomic segments, we denote the copy number mutation profile *j* by 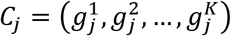. We can obtain the posterior probability that a given cell harbors copy number profile *C*_*j*_ by

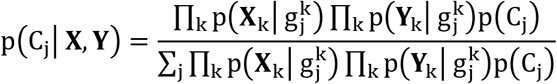

For example, the posterior probability that a cell is diploid in every region is p(C_0_), where *C*_0_ = ((1,1), (1,1), … , (1,1)). The posterior probability that a cell belongs to a specific clade (in particular, the tumor lineage) in the phylogeny is then equal to the sum of the probabilities that the cell harbors each of the possible genotypes included in the clade.

### WGS copy number analysis

We used hmftools^42^ to perform unmatched CNV analysis of the WGS data from the MM dataset. The COLBALT and AMBER modules were used to obtain the log read depth ratios (logR) and the BAF profiles, respectively. The PURPLE module was used to determine tumor ploidy and purity. We obtained continuous copy number segments based on the read depth ratios using the *pcf* function of *copynumber* R package^43^, with a gamma parameter of 12000. Significantly altered segments were determined by a threshold of logR > 0.25, logR < −0.25, BAF > 0.75 for amplifications, deletions, and CNLoH, respectively.

### Benchmarking the effect of population phasing prior on the detection of allelic imbalance

Using the TNBC4 dataset and cell annotations from the original paper, we created subsampled datasets composed of different tumor cell fractions. We defined the genomic boundaries of LoH events from the all-tumor pseudobulk. Using this setup, we performed three sets of benchmarking experiments. First, to benchmark the effect of phasing prior on the detection of subclonal allelic imbalance from heterogenous cell populations, we randomly sampled genomic segments with a fixed length (250 SNPs) from known LoH regions for each mixture proportion. We additionally sampled segments from the all-normal pseudobulks to serve as true negative examples. We then scored the allele profile of each sampled segment using the haplotype-naïve HMM and the haplotype-enhanced HMM. Using these scores, we calculated an AUC for each tumor-normal mixture proportion. Second, to benchmark the effect of phasing prior on allele classification (major vs minor haplotype) from mixture pseudobulks, we defined ground-truth haplotypes in known LoH regions using the observed BAFs in the all-tumor pseudobulk (BAF < 0.5, minor; BAF ≥ 0.5, major). We classified the alleles using the haplotype-naïve HMM and the haplotype-enhanced HMM for each mixture proportion. We then calculated the proportions of alleles correctly classified as a measure of model performance. Third, to benchmark the effect of phasing prior on single-cell event detection, we split the cells into training (70%) and testing sets (30%). We classified the alleles using the two models in known LoH regions using pseudobulks created from cells in the training set, and then used the obtained haplotypes to calculate allele posteriors of single cells in the test set. The existing tumor versus normal annotation was used as ground-truth labels for each cell and each event. We calculated an overall AUC (aggregating across events) for each tumor-normal mixture fraction.

### Benchmarking CNV detection accuracy

We evaluated the overall copy number profile reconstruction quality by Numbat and three other methods (CopyKAT, InferCNV, HoneyBADGER) using 5 MM samples (58408_Primary, 47491_Primary, 27522_Relapse_2, 37692_Primary, 59114_Relapse_1) with sample-matched flow-sorted WGS. Since Numbat, HoneyBADGER, and InferCNV identify CNVs from pseudobulks, we supplied the pseudobulk profile of all tumor cells. For CopyKAT, we summarized the consensus tumor copy number profile by averaging the copy number intensities for each genomic bin across all tumor cells. Since CopyKAT does not explicitly call copy number events, we applied a threshold of +0.03 and −0.03 to identify amplified and deleted segments. For HoneyBADGER, we took the union of events identified by the allele and expression approach. All tools were ran on default parameters and the HCA collection was used as diploid reference^44^. To evaluate the event detection performance, we computed precision and recall based on the aberrant/non-aberrant status of each marker using the DNA profile as ground truth. All types of events were considered (amplifications, deletions, CNLoH). To benchmark single-cell CNV testing accuracy, we first defined events from the DNA profile for each sample (event boundaries, event type, allele phasing). We did not include regions that appear to be affected by complex events (e.g. chr14 of 58408_Primary, Supplementary Figure 4) or subclonal events (e.g. chr16q deletion in 27522_Relapse_2, Supplementary Figure 4) as judged from the DNA profiles. We then computed a score of each event for each individual cell using the four different methods. For Numbat and HoneyBADGER, the event posterior probability was used as the score. For InferCNV and CopyKAT, we defined the score as the average smoothed expression intensity in the region affected by the event. Scores of CNLoH events is set to 0 for all allele-agnostic approaches. As an approximation of the cell-specific genotype ground truth, we assumed that the CNV events are present in all tumor cells and absent in all normal cells. For each event, we calculated an AUC based on the single-cell event scores from each method.

### Clonal decomposition using the Numbat iterative approach

We ran Numbat for two iterations on all datasets, with transition probability t = 10^−5^ for MM series and t = 10^−3^ for ATC/TNBC series to reflect the more complex genomic profile in solid tumors. For TNBC1, since shared diploid regions could not be identified, we manually supplied chromosomes 13,14,19 (containing CNLoH) as baseline when running Numbat. The HCA collection was used as the diploid reference for all datasets^44^.

### Benchmarking tumor versus non-malignant cell classification accuracy

We identified true tumor cells in the three datasets based on combined evidence of expression-based clustering, cell-type or tumor specific marker expression, and aneuploidy evidence. For the ATC and TNBC series, the tumor versus non-tumor labels from the original publication were used, and tumor-specific markers (*EPCAM* for TNBC, *KRT8* for ATC) were used as visual reference in Supplementary Figure 7–8. We excluded ATC5 from the benchmark due to the lack of clear expression of *KRT8*. For the MM series, we identified plasma cell clusters based on *MZB1* expression (Supplementary Figure 9). In one of the samples (27522_Relapse_2), both normal and malignant plasma cells are present, and the malignant plasma cell cluster was identified by upregulated *FGFR3* expression (due to t(4;14) translocation) as described in the original publication^39^. To evaluate performance, we calculated classification accuracy based on the ground-truth labels and the predictions made by the two methods. For Numbat, cells with combined aneuploidy probability > 0.5 are designated as tumor and normal otherwise. For CopyKAT, the tumor/normal predictions from the original paper was used for the TNBC and MDA series, and for the MM dataset, predictions were generated by running CopyKAT using its default parameters.

### Gene set enrichment analysis

We used the LIGER R package (https://github.com/JEFworks/liger) to perform the gene set enrichment analysis between clonal cell populations. Hallmark gene sets (n=50) were obtained from MSigDB. Only genes with at least one read count in at least 5 cells were used as input. We used the Holm-Bonferroni method to adjust for multiple comparisons. Significantly enriched gene sets were filtered by *Q* value < 0.05 and that the sign of the edge value is consistent with the enrichment direction (i.e. a positive enrichment is consistent with a positive edge value, and a negative enrichment is consistent with a negative edge value).

### Differential gene expression analysis

We used the Mann Whitney U test implemented in pagoda2 (https://github.com/kharchenkolab/pagoda2) to identify confident differentially expressed genes between subclones, treating each clone as a pseudobulk. We used the default parameter settings, with a Z score threshold of 3.

## Code availability

The Numbat algorithm is available at https://github.com/kharchenkolab/Numbat.

## Data availability

The scRNA-Seq and WGS validation data from the WASHU multiple myeloma study can be accessed through GEO under the accession GSE148673 and SRA under the accession PRJNA625321. The scRNA-Seq data from the MDA study can be accessed through GEO under the accession GSE148673 and SRA under the accession PRJNA694128. The HCA collection of reference expression profiles can be obtained from Synapse under the ID syn21041850.

## Author contributions

P.V.K. and T.G. formulated the study and the overall approach. A.K. carried out proof-of-concept tests of population-based phasing. T.G. developed the detailed algorithms with advice from P.V.K, R.A.S., H.S. and P.-R.L. T.G. implemented the Numbat package. T.G. and P.V.K. drafted the manuscript. All authors provided suggestions and corrections on the manuscript text.

---

**Supplementary Figure 1:**
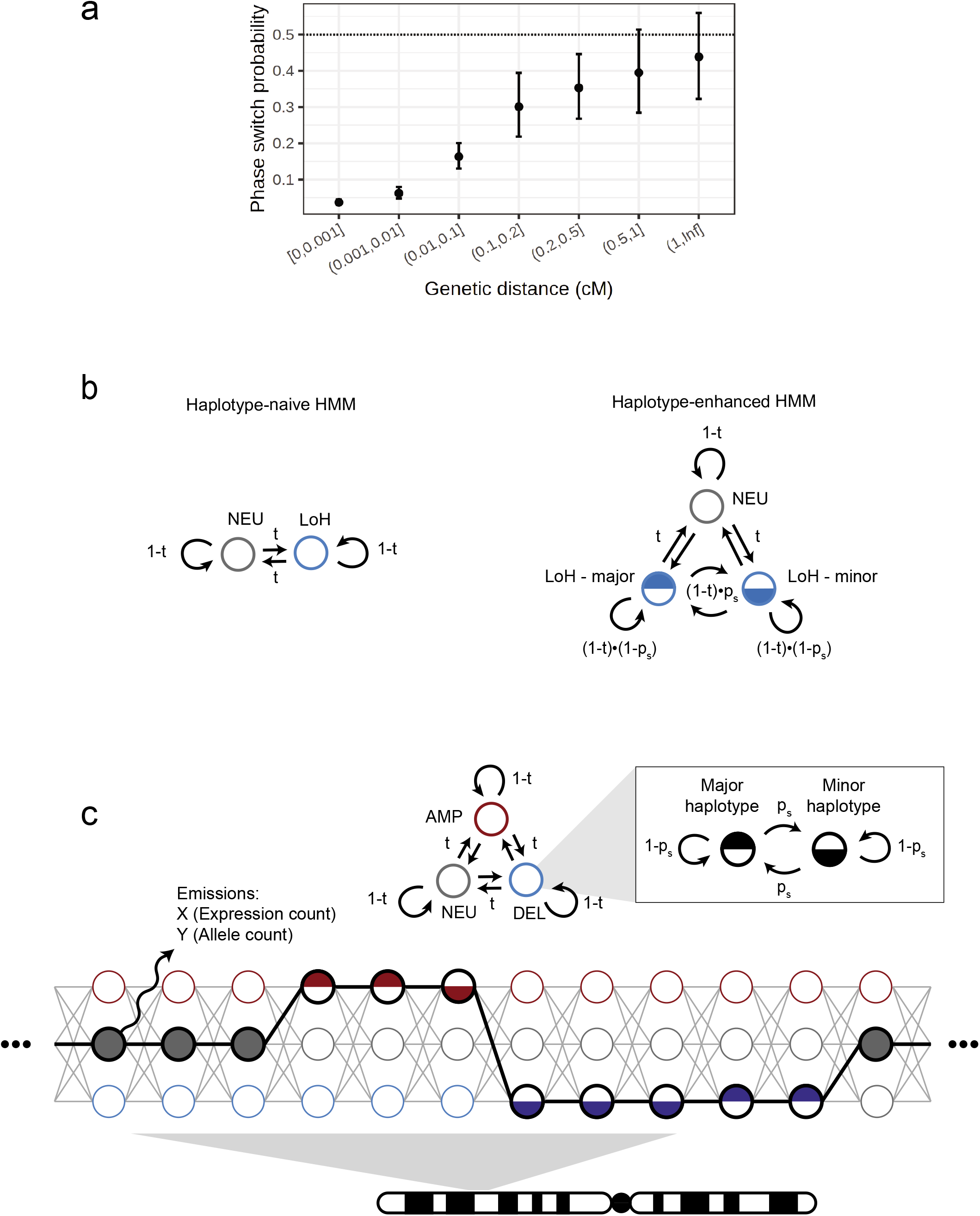
**(a)** Phase switch probability as a function of genetic distance, estimated from alleles phased from LoH regions in TNBC4. cM, centimorgan. Error bar represents 95% CI derived from a binomial test. **(b)** Schematic of conventional and haplotype-aware allele HMM. **(c)** Schematic of the Numbat joint HMM. Only three copy number states (neutral, deletion, amplification) and two haplotype states (major and minor) are included for illustrative purposes.

**Supplementary Figure 2:**
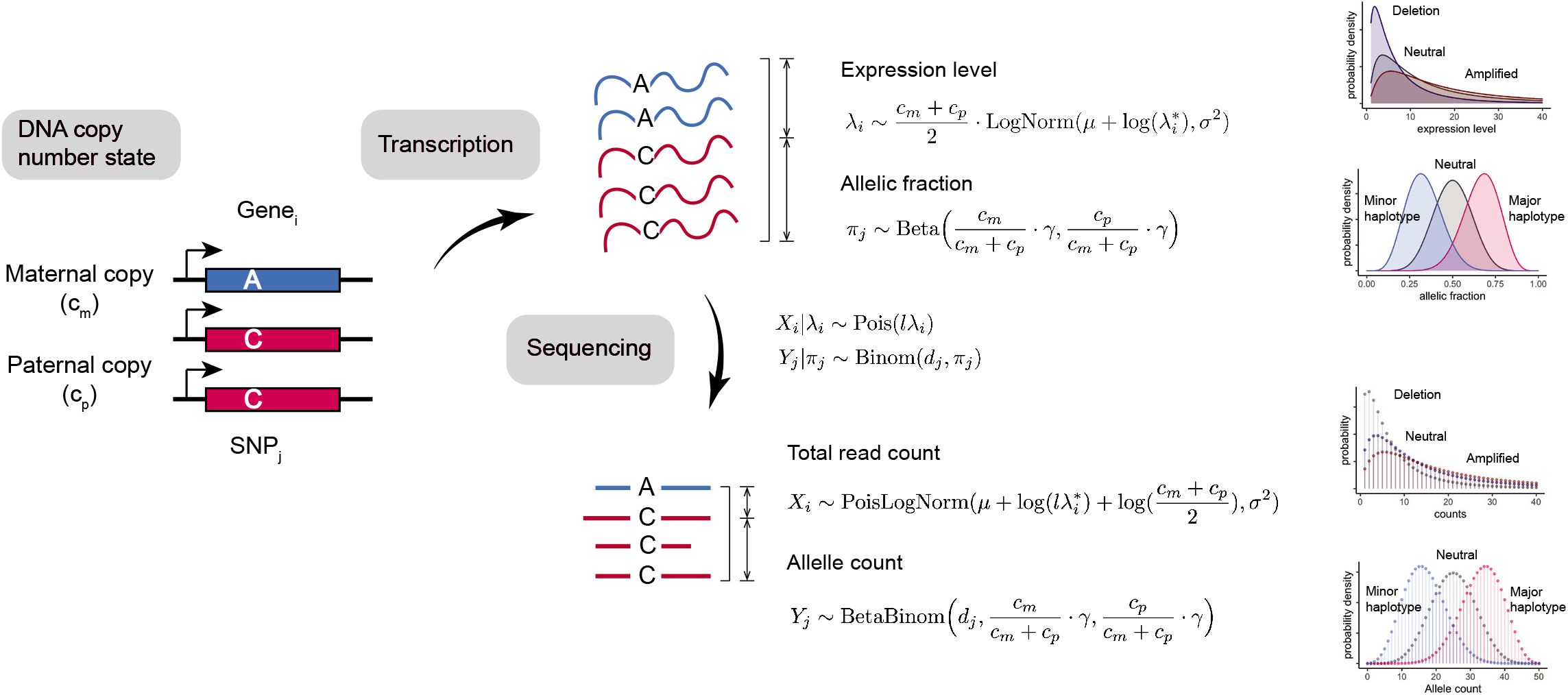
Generative model of gene expression and allele counts from scRNA-seq experiments.

**Supplementary Figure 3:**
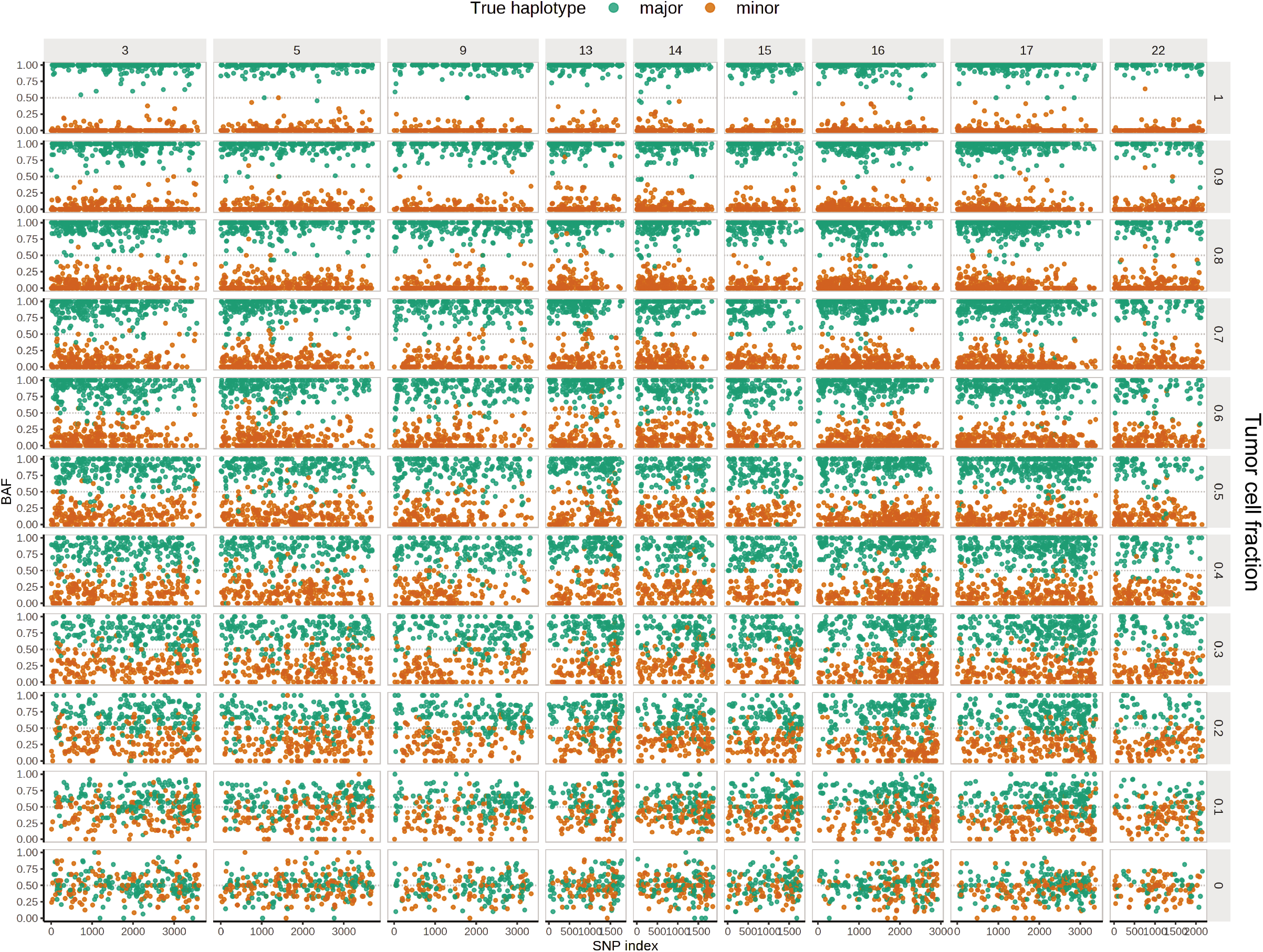
*In silico* serial dilution experiment of TNBC4 tumor and normal cells. Pseudobulk profiles were created from mixtures of various proportions of tumor and normal cells.

**Supplementary Figure 4:**
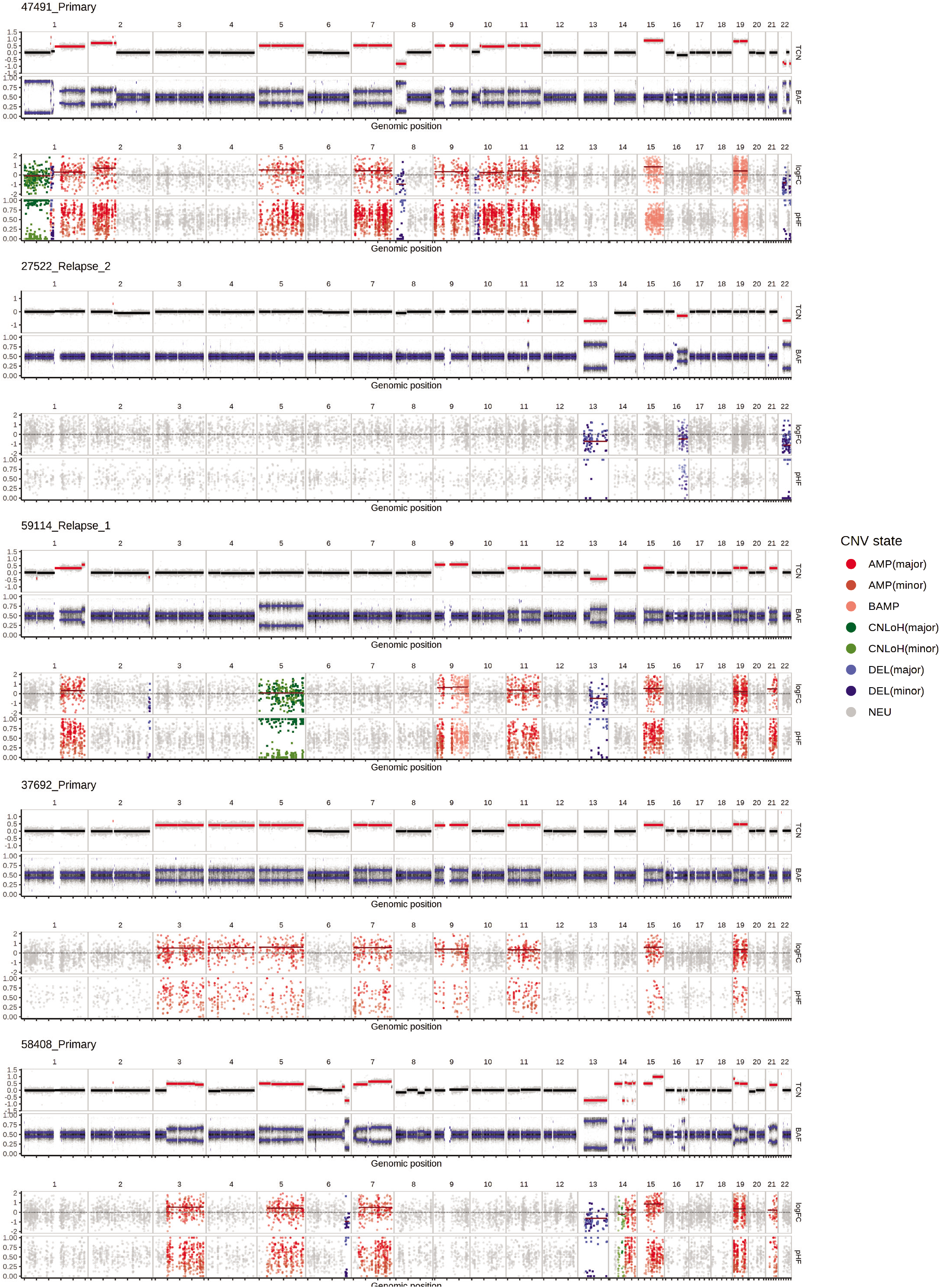
Numbat accurately detects copy number variations from scRNA-seq data. For each sample, the top panel shows the bulk DNA copy number profile obtained from flow-sorted WGS. The bottom panel shows the inferred copy number profile from scRNA data using Numbat. BAMP, balanced amplification. pHF, paternal haplotype frequency.

**Supplementary Figure 5:**
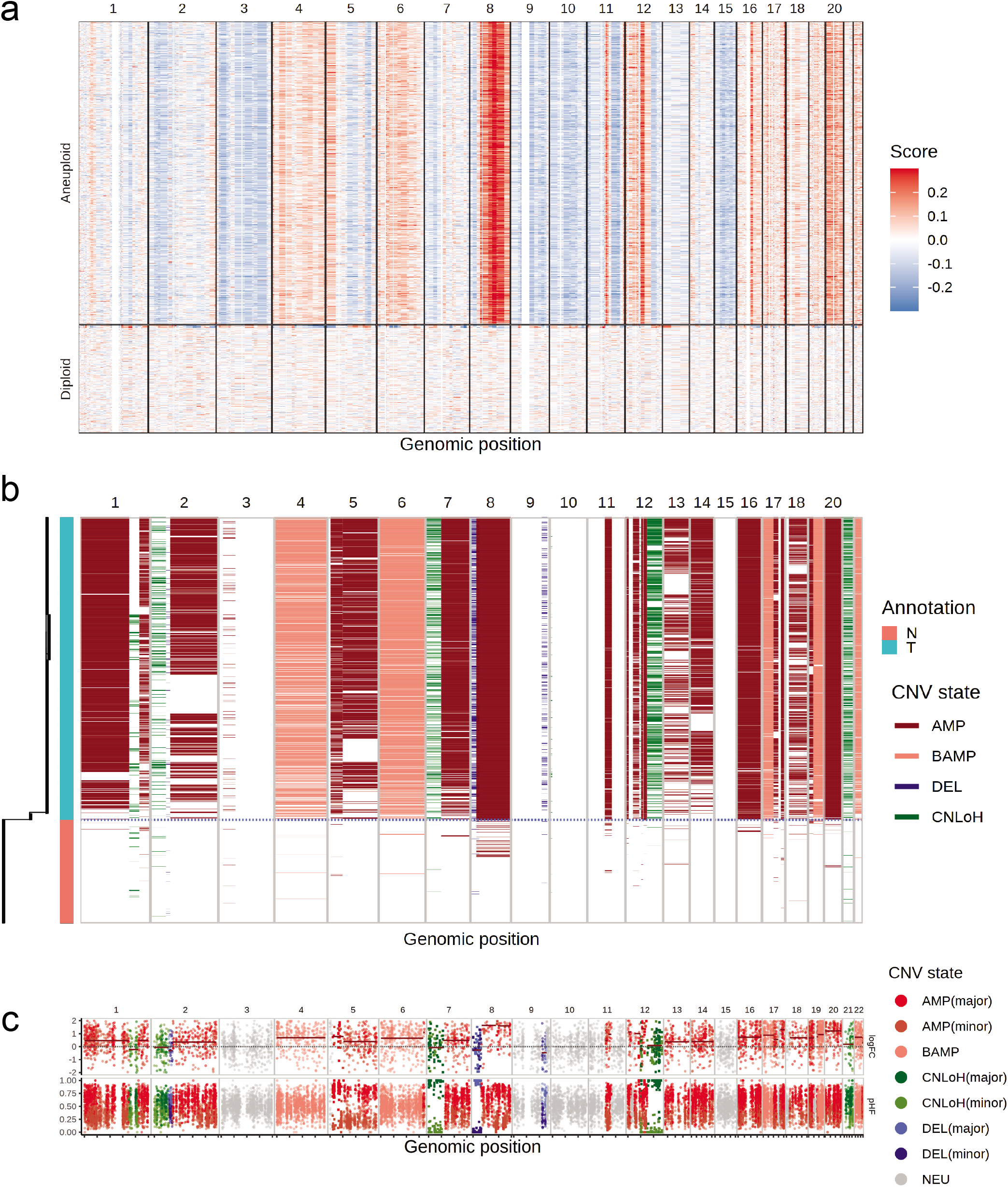
Numbat and CopyKAT analysis of DCIS1. **(a)** CopyKAT analysis of DCIS1. **(b)** Reconstructed single-cell copy number profile and clonal phylogeny by Numbat. The annotation corresponds to cell type ground truth (N, normal. T, tumor). **(c)** Numbat bulk CNV profile of tumor cells. BAMP, balanced amplification.

**Supplementary Figure 6:**
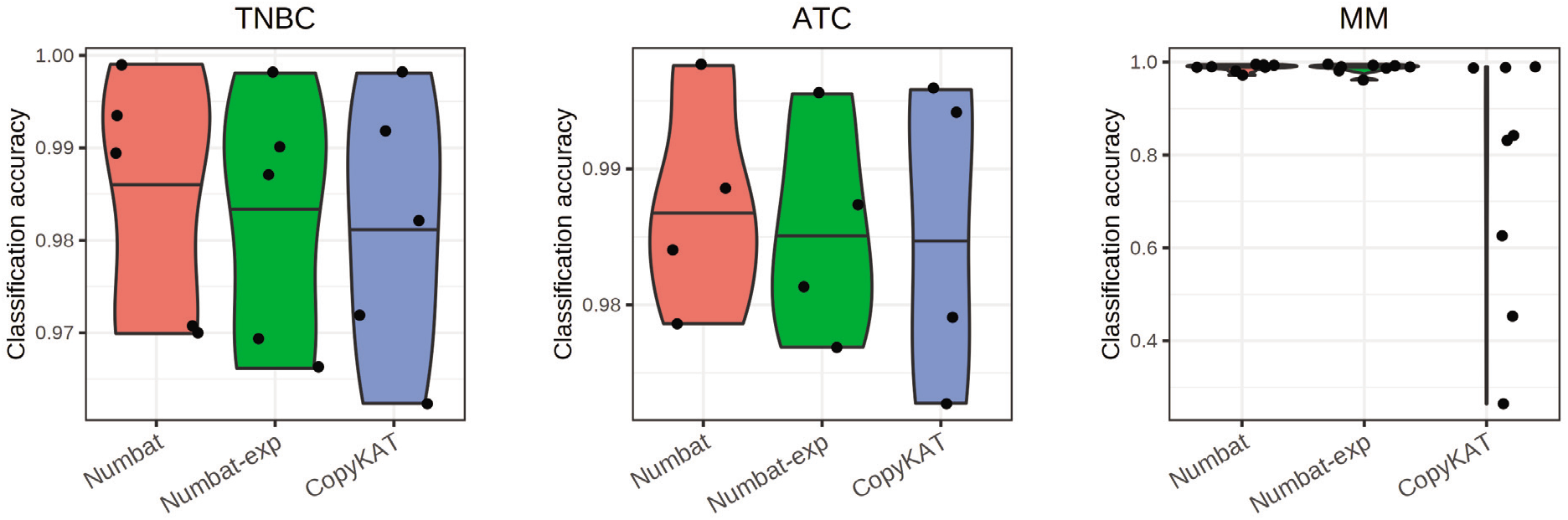
Tumor versus normal cell classification accuracy of Numbat joint model, Numbat expression-only model, and CopyKAT. Each dot represents a distinct sample (TNBC, n = 5; ATC, n = 4; MM, n = 8). Center line, median.

**Supplementary Figure 7:**
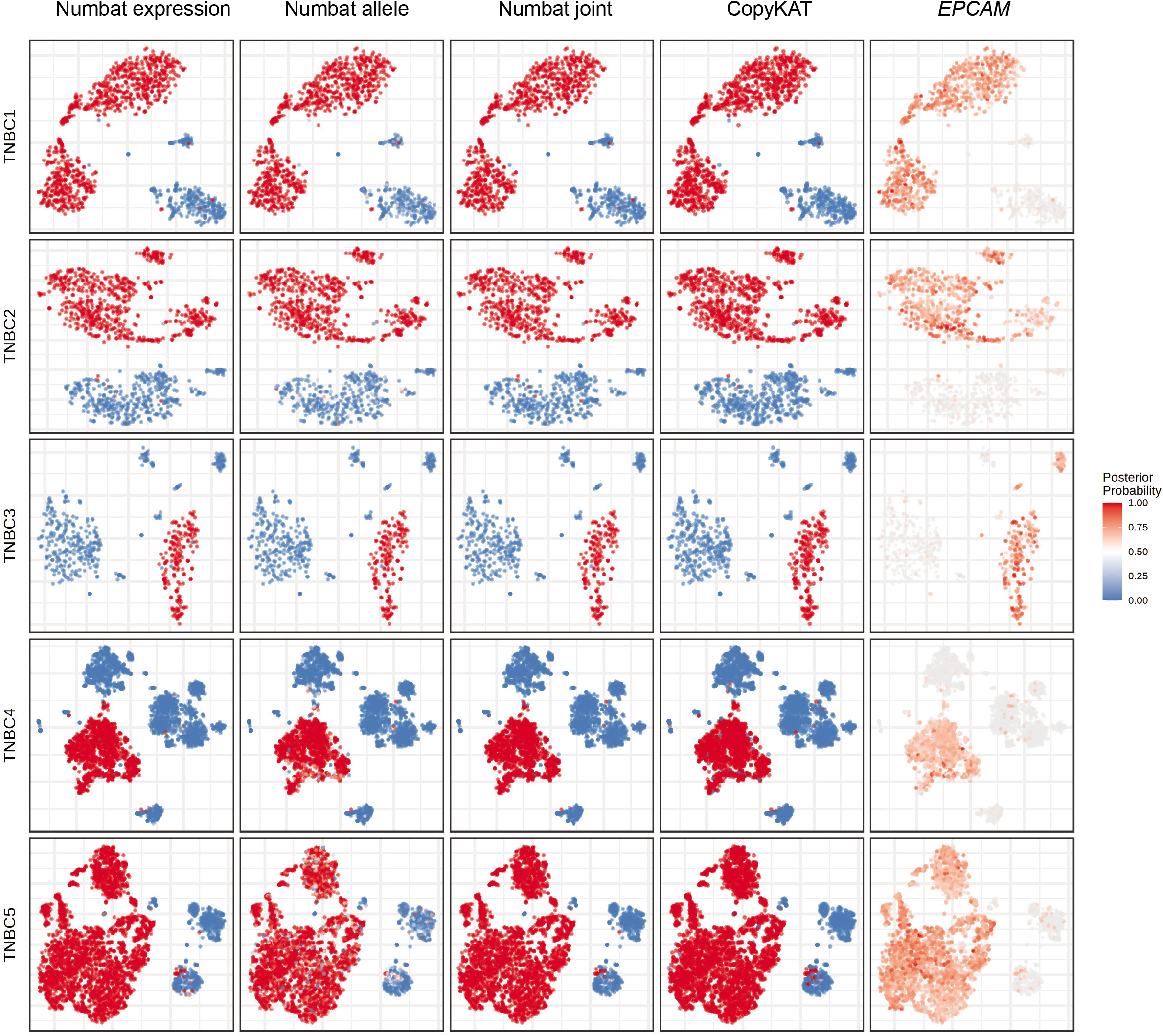
Numbat reliably distinguishes tumor and normal cells (TNBC series). Red, inferred tumor cells. Blue, inferred non-malignant cells. For each sample, the series of figures respectively show the aneuploidy probability by expression evidence, that by allele evidence, that by combined evidence, CopyKAT prediction, and tumor marker expression in a t-SNE embedding of gene expression profiles.

**Supplementary Figure 8:**
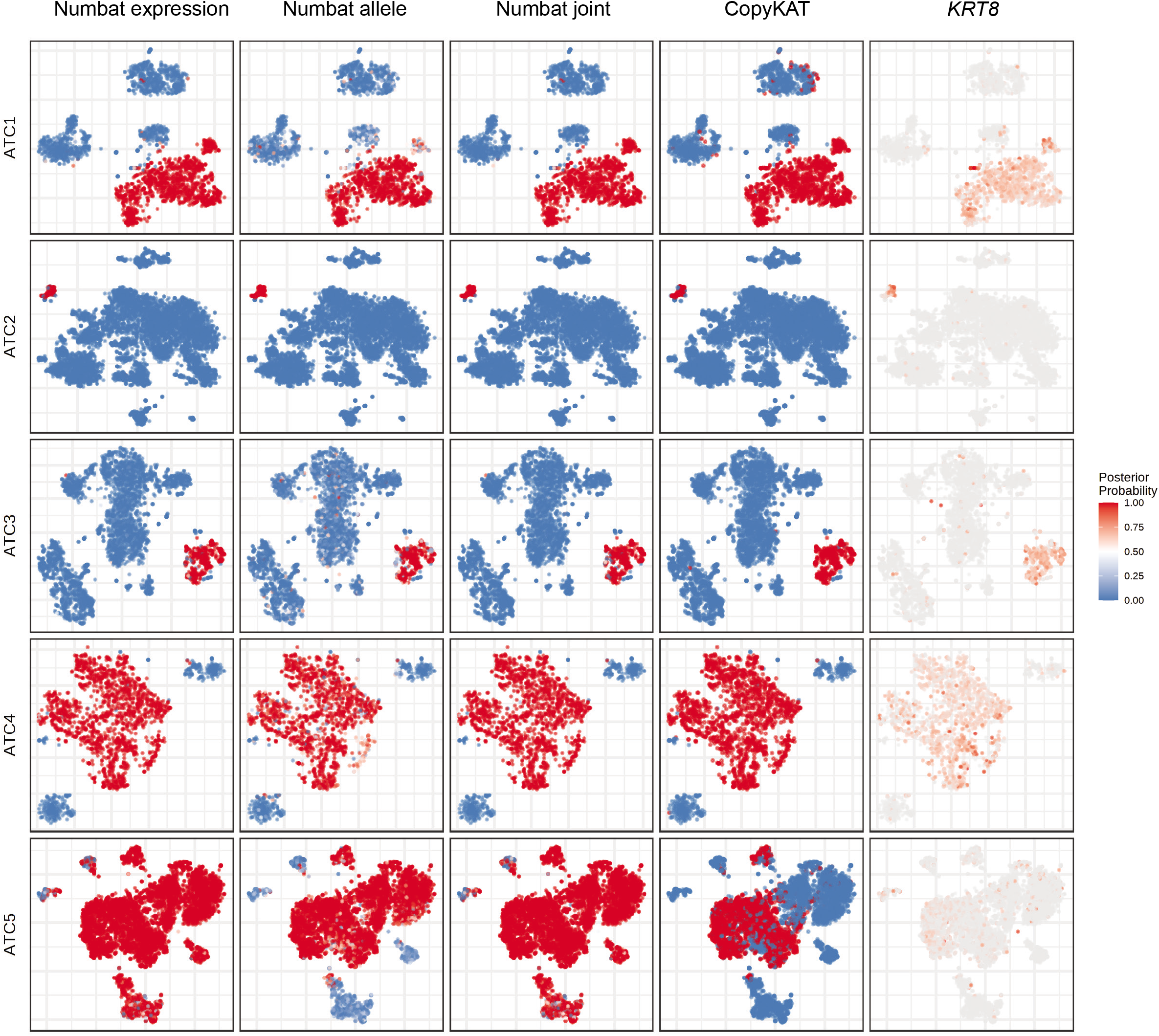
Numbat reliably distinguishes tumor and normal cells (ATC series). Red, inferred tumor cells. Blue, inferred non-malignant cells. For each sample, the series of figures respectively show the aneuploidy probability by expression evidence, that by allele evidence, that by combined evidence, CopyKAT prediction, and tumor marker expression in a t-SNE embedding of gene expression profiles. ATC5 was excluded from the benchmark due to lack of clear expression of *KRT8*.

**Supplementary Figure 9:**
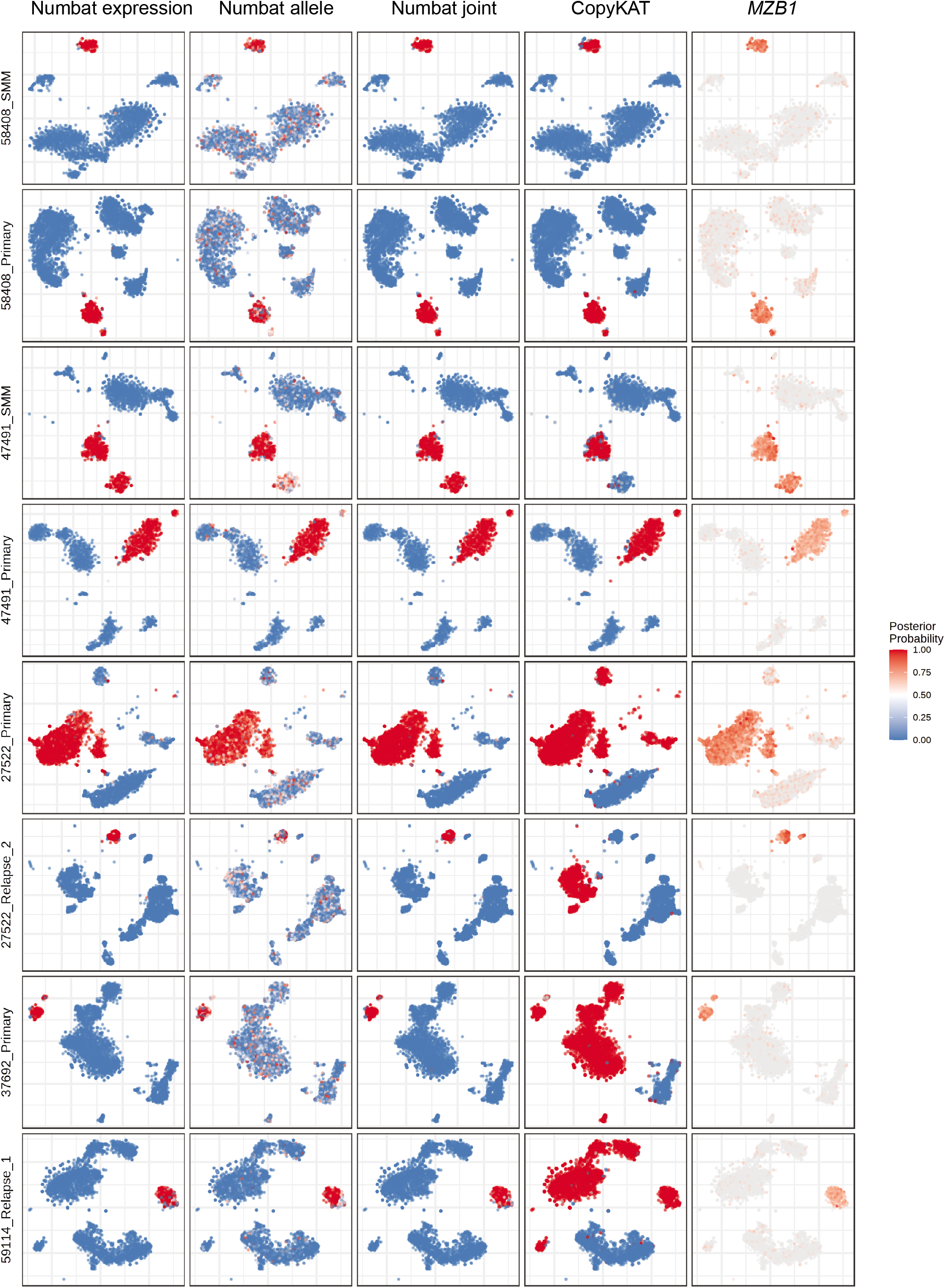
Numbat reliably distinguishes tumor and normal cells (MM series). Red, inferred tumor cells. Blue, inferred non-malignant cells. For each sample, the series of figures respectively show the aneuploidy probability by expression evidence, that by allele evidence, that by combined evidence, CopyKAT prediction, and tumor marker expression in a t-SNE embedding of gene expression profiles.

**Supplementary Figure 10:**
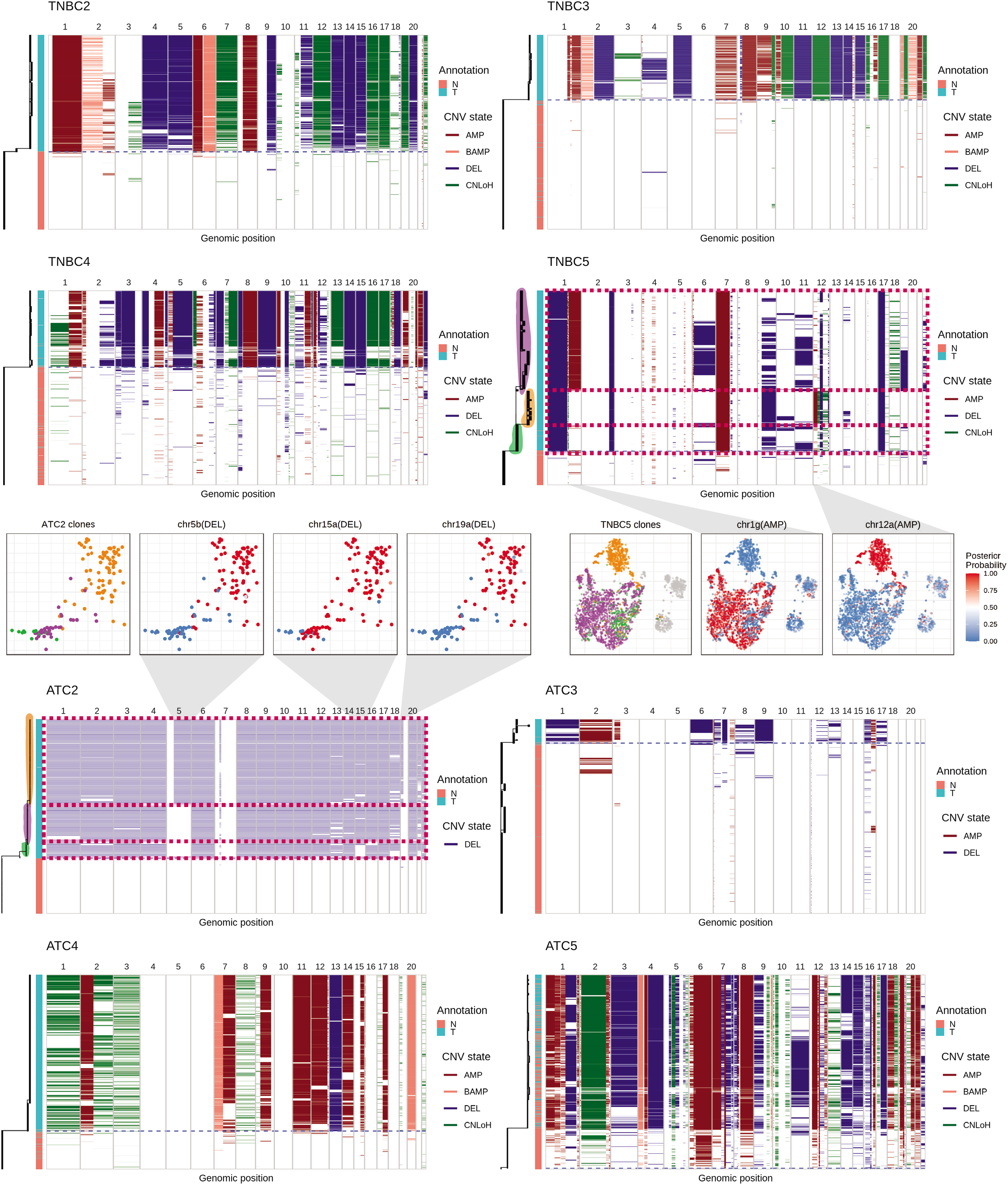
Single-cell copy number profile and phylogeny reconstructed by Numbat (TNBC and ATC). Branch lengths correspond to the number of mutations. Blue dashed line separates predicted tumor and normal cells. Confident subclones are highlighted and marked by red rectangles. The cell annotation from the original publication was used (T, tumor cells. N, normal cells). In this figure, normal cells from ATC2 were subsampled before running Numbat to better visualize the tumor subclonal phylogeny. In ATC5, some tumor cells were likely mis-annotated as normal in the original annotation.

**Supplementary Figure 11:**
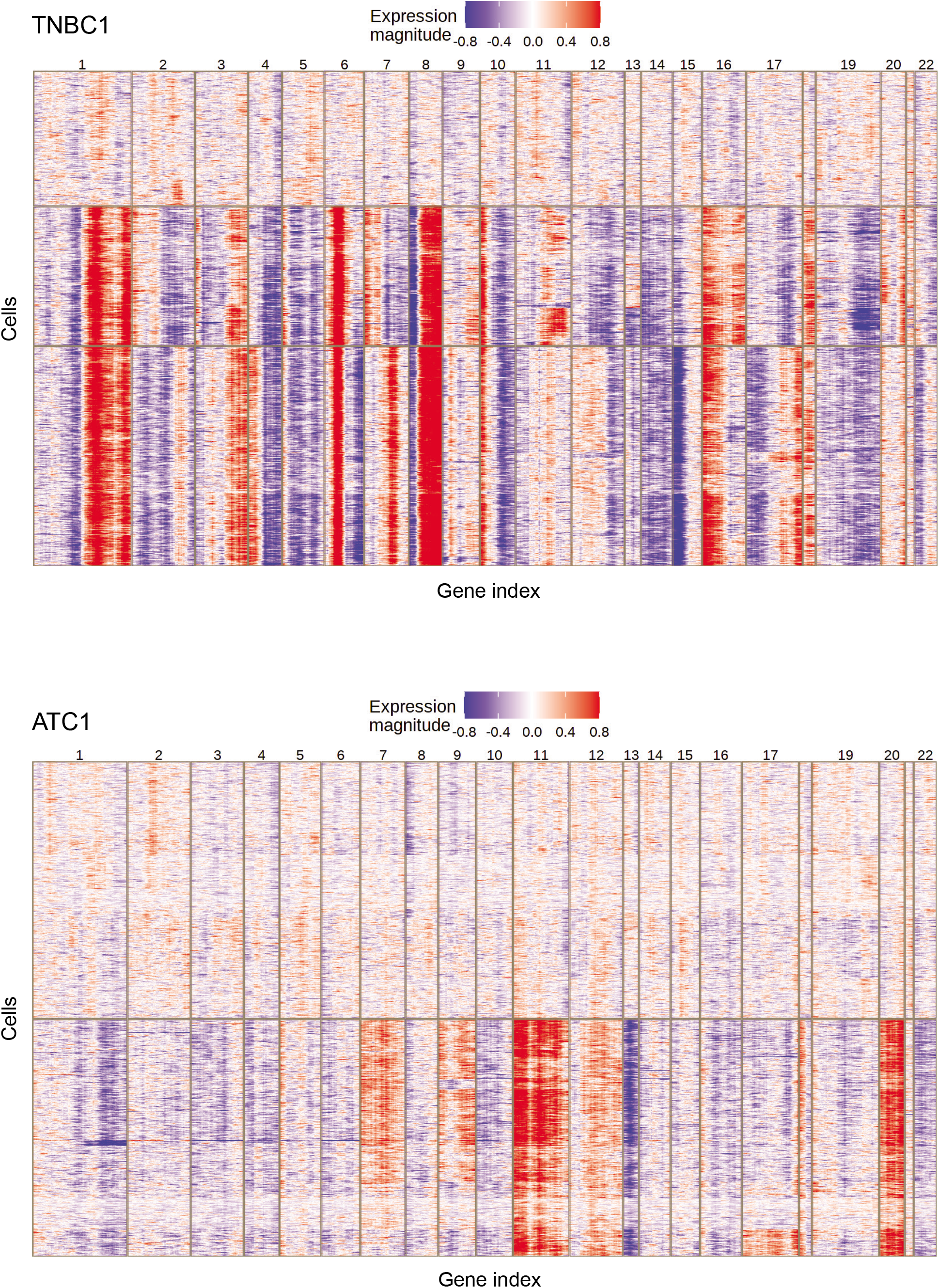
Window-smoothed expression profile and hierarchical clustering of TNBC1 and ATC1 cells.

**Supplementary Figure 12:**
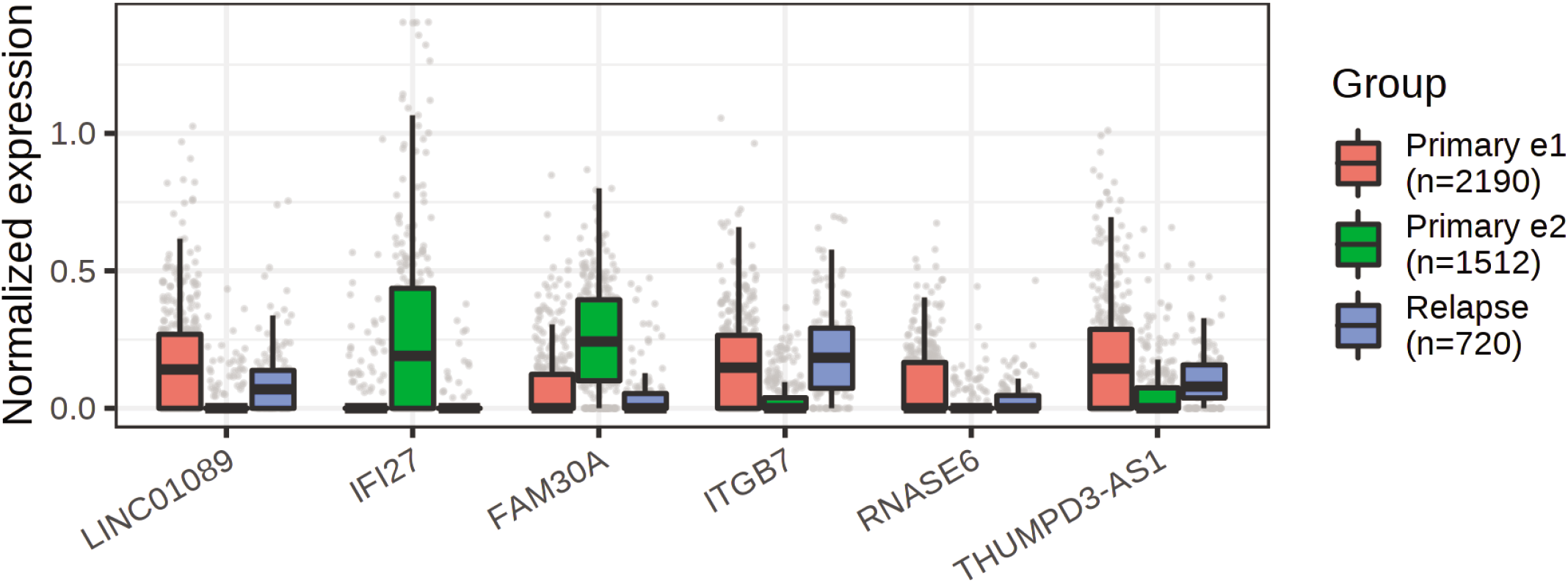
DE markers of expression clusters in multiple myeloma patient 27522. Each dot represents a distinct cell. Center line, median; box limits, upper and lower quartiles; whiskers, 1.5x interquartile range.

**Supplementary Table 1:** Sample information and sequencing characteristics of scRNA-seq datasets analyzed in the study.

| Sample ID | Patient ID | Cancer type | Number of cells | Protocol | Sorted WGS | Study |
| --- | --- | --- | --- | --- | --- | --- |
| 58408_SMM | 58408 | MM | 2426 | 10X 3' v2 | Yes | Liu et al. 2021 |
| 58408_Primary | 58408 | MM | 3890 | 10X 3' v2 | Yes | Liu et al. 2021 |
| 47491_SMM | 47491 | MM | 1769 | 10X 3' v2 | Low purity | Liu et al. 2021 |
| 47491_Primary | 47491 | MM | 1545 | 10X 3' v2 | Yes | Liu et al. 2021 |
| 37692_Primary | 37692 | MM | 3080 | 10X 3' v2 | Yes | Liu et al. 2021 |
| 59114_Relapse_1 | 59114 | MM | 3426 | 10X 3' v2 | Yes | Liu et al. 2021 |
| 27522_Primary | 27522 | MM | 4271 | 10X 3' v2 | No | Liu et al. 2021 |
| 27522_Remission | 27522 | MM | 727 | 10X 3' v2 | No | Liu et al. 2021 |
| 27522_Relapse_1 | 27522 | MM | 2280 | 10X 3' v2 | No | Liu et al. 2021 |
| 27522_Relapse_2 | 27522 | MM | 3499 | 10X 5' | Yes | Liu et al. 2021 |
| ATC1 | ATC1 | ATC | 2203 | 10X 3' v3 | No | Gao et al. 2021 |
| ATC2 | ATC2 | ATC | 6226 | 10X 3' v3 | No | Gao et al. 2021 |
| ATC3 | ATC3 | ATC | 3264 | 10X 3' v3 | No | Gao et al. 2021 |
| ATC4 | ATC4 | ATC | 1731 | 10X 3' v3 | No | Gao et al. 2021 |
| ATC5 | ATC5 | ATC | 6144 | 10X 3' v3 | No | Gao et al. 2021 |
| TNBC1 | TNBC1 | TNBC | 1097 | 10X 3' v2 | No | Gao et al. 2021 |
| TNBC2 | TNBC2 | TNBC | 1034 | 10X 3' v2 | No | Gao et al. 2021 |
| TNBC3 | TNBC3 | TNBC | 532 | 10X 3' v2 | No | Gao et al. 2021 |
| TNBC4 | TNBC4 | TNBC | 3056 | 10X 3' v3 | No | Gao et al. 2021 |
| TNBC5 | TNBC5 | TNBC | 3225 | 10X 3' v3 | No | Gao et al. 2021 |
| DCIS1 | DCIS1 | DCIS | 1480 | 10X 3' v2 | No | Gao et al. 2021 |

**Supplementary Table 2:**
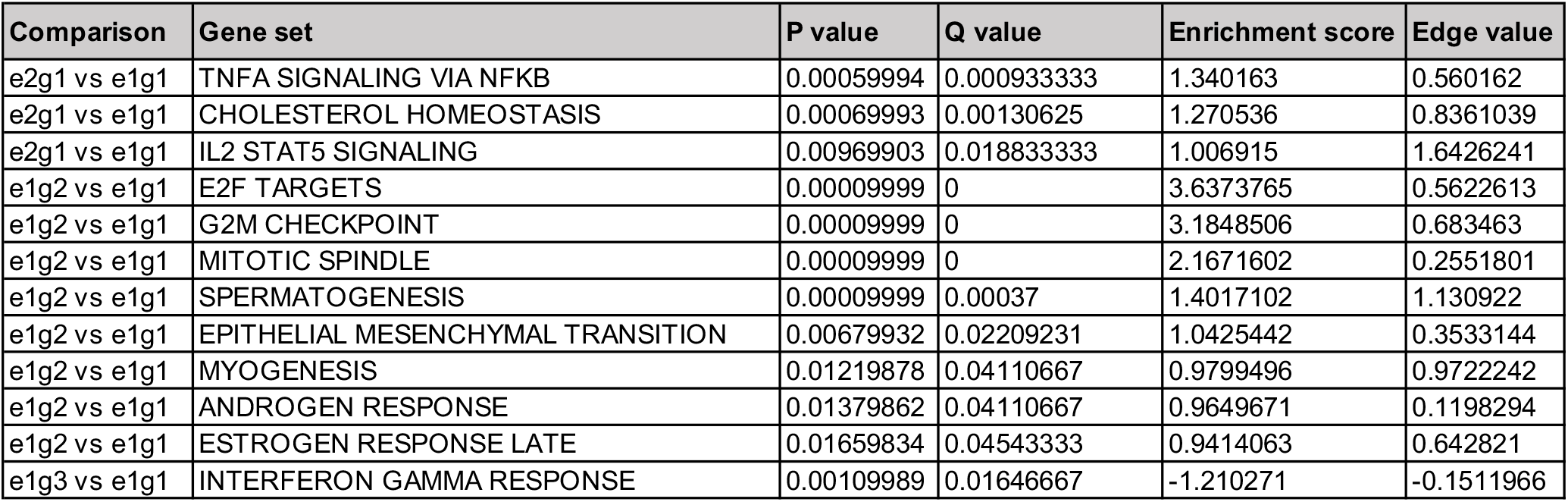
List of significantly enriched pathways in multiple myeloma patient 27522.

| Comparison | Gene set | P value | Q value | Enrichment score | Edge value |
| --- | --- | --- | --- | --- | --- |
| e2g1 vs e1g1 | TNFA SIGNALING VIA NFKB | 0.00059994 | 0.000933333 | 1.340163 | 0.560162 |
| e2g1 vs e1g1 | CHOLESTEROL HOMEOSTASIS | 0.00069993 | 0.00130625 | 1.270536 | 0.8361039 |
| e2g1 vs e1g1 | IL2 STAT5 SIGNALING | 0.00969903 | 0.018833333 | 1.006915 | 1.6426241 |
| e1g2 vs e1g1 | E2F TARGETS | 0.00009999 | 0 | 3.6373765 | 0.5622613 |
| e1g2 vs e1g1 | G2M CHECKPOINT | 0.00009999 | 0 | 3.1848506 | 0.683463 |
| e1g2 vs e1g1 | MITOTIC SPINDLE | 0.00009999 | 0 | 2.1671602 | 0.2551801 |
| e1g2 vs e1g1 | SPERMATOGENESIS | 0.00009999 | 0.00037 | 1.4017102 | 1.130922 |
| e1g2 vs e1g1 | EPITHELIAL MESENCHYMAL TRANSITION | 0.00679932 | 0.02209231 | 1.0425442 | 0.3533144 |
| e1g2 vs e1g1 | MYOGENESIS | 0.01219878 | 0.04110667 | 0.9799496 | 0.9722242 |
| e1g2 vs e1g1 | ANDROGEN RESPONSE | 0.01379862 | 0.04110667 | 0.9649671 | 0.1198294 |
| e1g2 vs e1g1 | ESTROGEN RESPONSE LATE | 0.01659834 | 0.04543333 | 0.9414063 | 0.642821 |
| e1g3 vs e1g1 | INTERFERON GAMMA RESPONSE | 0.00109989 | 0.01646667 | -1.210271 | -0.1511966 |

